# Copy number variants in clinical WGS: deployment and interpretation for rare and undiagnosed disease

**DOI:** 10.1101/245100

**Authors:** Andrew M Gross, Subramanian S. Ajay, Vani Rajan, Carolyn Brown, Krista Bluske, Nicole Burns, Aditi Chawla, Alison J Coffey, Alka Malhotra, Alicia Scocchia, Erin Thorpe, Natasa Dzidic, Karine Hovanes, Trilochan Sahoo, Egor Dolzhenko, Bryan Lajoie, Amirah Khouzam, Shimul Chowdhury, John Belmont, Eric Roller, Sergii Ivakhno, Stephen Tanner, Julia McEachern, Tina Hambuch, Michael Eberle, R Tanner Hagelstrom, David R Bentley, Denise L Perry, Ryan J Taft

## Abstract

**Purpose:** Current diagnostic testing for genetic disorders involves serial use of specialized assays spanning multiple technologies. In principle, whole genome sequencing (WGS) has the potential to detect all genomic mutation types on a single platform and workflow. Here we sought to evaluate copy number variant (CNV) calling as part of a clinically accredited WGS test.

**Methods:** Using a depth-based copy number caller we performed analytical validation of CNV calling on a reference panel of 17 samples, compared the sensitivity of WGS-based variants to those from a clinical microarray, and set a bound on precision using orthogonal technologies. We developed a protocol for family-based analysis, annotation, filtering, visualization of WGS based CNV calls, and deployed this across a clinical cohort of 79 rare and undiagnosed cases.

**Results:** We found that CNV calls from WGS are at least as sensitive as those from microarrays, while only creating a modest increase in the number of variants interpreted (~10 CNVs per case). We identified clinically significant CNVs in 15% of the first 79 cases analyzed. This pipeline also enabled identification of cases of uniparental disomy (UPD) and a 50% mosaic trisomy 14. Directed analysis of some CNVs enabled break-point level resolution of genomic rearrangements and phasing of *de-novo* CNVs.

**Conclusion:** Robust identification of CNVs by WGS is possible within a clinical testing environment, and further developments will bring improvements in resolution of smaller and more complex CNVs.

## Introduction

Variation in DNA copy-number is a well-described cause of human genetic disease (Lupski, 2015). Copy-number variants (CNV) associated with human pathologies range from chromosomal aneuploidy, to micro-duplication and -deletion syndromes, and include smaller structural variants that affect single genes and exons (Harel and Lupski, 2017; Lupski, 2015; Riggs et al., 2012; Swaminathan et al., 2012; Vulto-van Silfhout et al., 2013). Karyotype and microarray have served as gold-standards in molecular diagnostics for CNVs, but the increasing number and complexity of possible genomic changes requires testing that can simultaneously address the complete range of cytogenetic abnormalities and smaller structural variants.

Whole genome sequencing (WGS) can be used to detect almost all classes of alleles. It is sensitive and specific for SNVs and indels (Kim et al., 2017; Van der Auwera et al., 2013), enables detection of complex repeat expansions (Dolzhenko et al., 2016) and proof-of-principle studies have shown the ability to detect copy-number <u>(Abyzov et al., 2011; Roller et al., 2016)</u> and structural variation (Sudmant et al., 2015). Approaches have been developed to enable CNV detection using other next generation sequencing (NGS) panels and exomes, which have improved diagnostic yield (Eisenberger et al., 2013; Tian et al., 2015), but have technical limitations arising from non-uniform sequencing depth, PCR artifacts, GC bias, and a larger variance in allele fraction (Lelieveld et al., 2015; Linderman et al., 2014; Meienberg et al., 2016; Meynert et al., 2014). In contrast, WGS sequencing depth is predictable and robust throughout the genome (Lionel et al., 2017; Meienberg et al., 2016), and eliminates the hazard of capture and PCR-based artifacts. This uniformity of signal enables sample-specific depth normalization that eliminates the need for batch processing (Boeva et al., 2012; Roller et al., 2016). Furthermore, coverage of the non-coding genome (including deep intronic regions) allows for increased resolution to detect small CNVs, more accurate estimation of variant boundaries, and in many cases direct evidence for the underlying DNA rearrangement via observation of paired-sequencing read alignments (Newman et al., 2015; Sudmant et al., 2015).

Here we describe the deployment of a CNV detection pipeline as a component of a clinical WGS (cWGS) diagnostic test for patients with rare or undiagnosed genetic disease (RUGD). As a single assay, WGS has the potential to benefit RUGD patients by enabling detection of multiple variant types simultaneously, decreasing the number of molecular tests performed, increasing the range of detectable disorders, and shortening the diagnostic odyssey. Below we describe the technical feasibility assessment and validation of WGS-based CNV calls compared to a microarray-based clinical diagnostic test, and the deployment of CNV calling as part of a cWGS test for RUGD.

## Methods

### CNV truthset generation and sensitivity assessment

Twenty reference samples (Coriell, Camden, NJ) events were chosen for validation (**Table S1**). Among these, 18 samples had known pathogenic CNVs representative of a large size range and inclusive of deletions and copy-number gain, and two samples were included as negative controls. Prior to sequencing and analysis, coordinates for ‘truth-set’ CNVs were compiled from descriptions on the Coriell website, reference publications or previously conducted microarray-based CNV analyses (**Table S1**, Tang et al). We note that while all cell-lines contain pathogenic CNVs which established the baseline for our sensitivity analysis, we also examined other all other CNVs detected in these samples by either microarray or WGS.

DNA samples were procured from Coriell and libraries were prepared using the Illumina TruSeq PCR-free kit and sequenced on HiSeq X with paired-end 150bp reads in the Illumina Clinical Services Laboratory (Illumina Inc, San Diego CA). Data were mapped to the hg19 reference genome with the ISAAC aligner (Raczy et al., 2013). The resulting BAM files were analyzed with the Canvas CNV caller (Roller et al., 2016, see also “Copy number variant detection and filtering”). In parallel, samples were assessed by an external clinical microarray lab (CombiMatrix Diagnostics, Irvine, CA), which included profiling on an Illumina 850k feature SNP array followed by automated CNV calling and manual curation by trained cytogeneticists. One sample failed microarray analysis, resulting in 17 positive control samples for further analysis.

To assess sensitivity, WGS and microarray call sets were compared (requiring 50% or 75% overlap) with reference calls (**Table 1**, **Table S2**, see **Supplemental Note**). For false-negative calls or calls with only partial overlap with the reference call, visualization of depth and microarray data was conducted to assess the accuracy of the call boundaries or identify discrepancies of WGS based boundaries with the vendor-supplied CNV annotation (**Results** and **Supplemental Note**).

**Table 1.** Summary of sensitivity of cWGS and clinical microarrays to annotated CNVs in cell-lines.

| event | size | Coriell Events | Called by Array* | Called by WGS* |
| --- | --- | --- | --- | --- |
| LOSS | 10kb-50kb | 5 | 3 (+1) | 4 |
|  | 50kb-100kb | 1 | 1 | 1 |
|  | 100kb-500kb | 9 | 3 (+1) | 6 |
|  | >500kb | 6 | 6 | 6 |
|  | Overall | 21 | 13 (61%) | 17 (80%) |
| GAIN | 10kb-50kb | 3 | 0 | 3 |
|  | 100kb-500kb | 5 | 4 | 4 |
|  | >500kb | 7 | 6 | 7 |
|  | Overall | 15 | 10 (67%) | 14 (93%) |
| All CNV calls |  | n=36 | 23 (64%) | 31 (86%) |
| * 50% overlap |  |  |  |  |
| Note that +1 indicates calls that were not in call set, but recovered in manual review |  |  |  |  |

### Assessment of cWGS CNV calling false positive rate

To determine an upper bound on the false positive rate for the WGS Canvas CNV calling pipeline, we systematically assessed CNV calls made on Platinum Genome NA12878 (Eberle et al., 2017) using the 1000 Genomes Project NA12878 reference assembled using a combination of Pacific Biosciences (PacBio) and BioNano DNA data (Pendleton et al., 2015). A CNV call was considered validated if there was at least 75% overlap with BioNano call boundaries. Calls with partial overlap were manually curated to assess possible false positive or partially called BioNano CNVs resulting in a low overlap (see **Supplemental Note**). CNVs that were not called in the PacBio+BioNano dataset were manually reviewed for the presence of discordant sequencing reads spanning the boundaries of a deletion or copy-number gain (indicative of a tandem-duplication), and the presence of hemizygous and homozygous deletions with similar breakpoints in an independent set of samples from population controls (N=3000).

### Clinical cohort inclusion criteria

Variant calling and interpretation of CNVs was deployed into the Illumina Clinical Services Lab (ICSL) as part of routine practice for RUGD cases finishing sequencing and primary analysis on or after June 2, 2016. Clinically relevant losses or gains greater than 10kb were reported. Since that time, 79 patients were consented for the TruGenome Undiagnosed Disease test to be performed by ICSL. These patients had a wide spectrum of phenotypes, as well as previous testing ranging from no prior molecular investigations to panels and whole exome sequencing. The age at the time of testing ranged from 1 year to 20 years.

### Copy number variant detection and filtering

CNV detection and annotation were validated and deployed using paired-end 150 nucleotide HiSeqX sequencing runs, processed through the Illumina’s secondary analysis pipeline for short-read alignment and variant calling.

CNVs are called from the generated BAM files using version 1.3.9 of the Canvas caller (Roller et al., 2016) under its germline WGS setting, with modifications to the default calling parameters as follows:

- The circular binary segmentation (CBS) segmentation algorithm is used as opposed to the Haar wavelet based default in Canvas v1.3.9. This is specified as a parameter on the Canvas command line invocation.
- In practice we often see fragmented large CNV events. To limit this, candidate CNV calls spaced by less than 100kb are merged into a single call. When such a merge occurs, the magnitude of the gap between segments and any implications on variant interpretation are assessed during manual curation.
- Support thresholds for candidate CNVs were dropped to 8 depth bins to increase sensitivity in 10-50 kB range (a depth bin is defined as a sequence range with an expected 100 reads mapping).
- An automated b-allele based ploidy correction step was omitted, to limit false negatives. Screening for presence of heterozygous variants in a candidate deletion was deferred to the manual curation stage.
- To include common variation, the grey list of filtered regions supplied within Canvas was omitted and replaced with a minimal list of chromosomal segments covering centromeres.
- Canvas quality scores were not used as a filter for candidate CNV events.

All CNV calls are processed through a series of automated filtering steps (**Figure S1**) to reduce false positives and limit downstream CNV curation (**Figure S3**) to those likely to have medical relevance.

#### Canvas grey-list filter

Canvas provides a set of grey-list regions that contains problematic genomic segments as well as common CNVs. In filtering, CNVs that have greater than 50% of their range spanned by grey list regions are filtered out.

#### Gene annotation and filtering

CNVs are annotated with overlapping or nearby (<5kb away) genes using RefSeq gene definitions. Calls with no gene annotation are filtered.

#### Population frequency annotation and filtering

CNV population frequency is estimated using an internal database of samples sequenced in the Illumina services lab and individually normalized through the Canvas CNV calling pipeline. Binned sequencing depth data (an intermediate output of Canvas) is mapped to a fixed 300bp uniform coordinate system to allow for efficient storage and recovery of data across many samples.

Due to uncertainty in the boundaries of many CNV calls, a heuristic calculation of CNV population frequency is implemented that includes (1) interrogation of the aggregate sequencing depth data across 3,000 genomes for the genomic interval defined by the CNV boundaries; (2) mean depth analysis for each sample compared to predefined thresholds calculated from the population given the expected ploidy of the region; and (3) the fraction of the population samples consistent with the proband GAIN or LOSS status is calculated. Note that for events on a sex chromosome, only samples with the same gender from the population are queried, and that this does not account for the magnitude of the copy number change, but rather only the direction of the change from the diploid or haploid expectation.

For interpretation, CNVs with a population frequency higher than 10% (~5% allele frequency) are filtered out, and the vast majority of CNV calls in the 1-10% range are not classified as being clinically significant after review (**Figure S2**).

### Interrogation of clinically relevant CNVs

When a CNV is annotated as of clinical interest by the curation process, additional bioinformatic analysis may be conducted to provide further annotation in order to aid in variant interpretation.

#### CNV Phasing

Where possible, the parental phasing is assessed by genotyping parental haplotypes using depth information, or using inheritance patterns of small variants when no evidence of a depth change is present in a parent (e.g. de-novo CNVs or duo cases where there is no evidence of a CNV in the sequenced parent).

The de-novo CNV phasing algorithm first constructs prior state probabilities given the genotypes of the parents and the known copy-number of proband. Given the prior probabilities of each transition, we compute the model likelihood for all possible inheritance assumptions, and the most likely model is selected.

For example, at a haploid (copy number 1) site where the mother is heterozygous (0/1) and the father is homozygous reference (0/0) for a given SNV:

- under the assumption that the CNV is inherited from the father, the probability of a REF or haploid SNV call in the proband are both 50%
- under the assumption that the CNV is inherited from the mother, the probability of a REF or haploid SNV call in the proband are 100% and 0% respectively

Probabilities for all inherence assumptions are calculated across all SNVs within the target region are calculated and the model is selected via a maximum likelihood criteria. For details of inheritance models and examples, see **Supplemental Note**.

#### Interrogation of structural variation at CNV boundaries

Sequencing reads adjacent to CNVs can provide evidence of complex chromosomal rearrangements. For CNVs indicative of large structural variants - including terminal chromosomal deletions, large tandem duplications and break-ends spanning non-homologous chromosomes - the Manta structural variant caller (Chen et al., 2016, version 0.29.3) was employed for further investigation. This enabled breakpoint linkage across multiple CNVs, and provided evidence for insertion of duplicated sequence into a chromosome. Additionally, reassembled breakpoints were visualized via realignment of sequencing reads using the SVViz program (Spies et al., 2015).

## Results

### Validation of the CNV calling pipeline

Assessment of 17 reference samples with known pathogenic CNVs (**Table S1**) showed that cWGS had greater sensitivity to detect known CNVs compared to microarrays (86% versus 64%, **Methods, Table 1, Table S2**) with the largest difference in smaller (<50kb) events (**Table 1**). For the five ‘truth set’ CNVs not recovered by cWGS, manual inspection of sequencing and genotyping arrays did not support a CNV in these regions (**Methods, Supplemental Note**).

To assess the cWGS false positive rate, CNVs were called on the 1000 Genomes Project sample NA12878 and assessed against an orthogonal technology genome assembly (Sudmant et al., 2015). Among 93 deletions, 48 had analogs in a dataset derived from long-read sequencing technology (Pendleton et al., 2015) (BioNano calls, **Methods**), and nine additional calls were supported by the presence of discordant sequencing reads and/or evidence of Mendelian inheritance across a population of samples. Thirty-nine percent (36/93) of the cWGS NA12878 CNV calls do not have support from orthogonal data or independent bioinformatics analysis, which will include both false positives and suspected true calls without external support. Because of limitations in all predicate CNV calling methods, the discrepancies may arise from either WGS or the alternative platforms.

We found that the majority of putative cWGS false positives can be addressed with minimal heuristic filters. Specifically, application of a size filter restricting CNVs >10kb, removal of CNVs in regions that have variable data quality, putative mosaics, and eliminating CNVs found in >10% of the population (drawn from a cohort of more than 3000 genomes, see **Methods**, **Figures 1-3**) reduced our CNV interpretation burden to an average of 11 calls per case (range 2-26). Given these findings, these heuristics were deployed as a component of the clinical WGS pipeline.

### Deployment of the CNV pipeline into the clinical lab

Seventy nine clinical cases were processed through the validated cWGS CNV pipeline between June 2, 2016 and April 19, 2017 and subjected to automated quality control, filtering, annotation, and visualization (**Figures S1–S3**, **Methods**). CNVs were curated and classified following the guidelines of the American College of Medical Genetics and Genomics (ACMG) ((Kearney et al., 2011; South et al., 2013), **Methods**, **Figure S2a-d**). After filtering based on internal allele frequency, on average, we reported 3 benign, 3 VUS-likely benign, and 4 VUS CNVs per case (**Figure S2e-g**). In 15% (11/79) of cases, we reported variants with pathogenic or “uncertain significance - likely pathogenic” classifications across a diverse set of patient phenotypes (**Table 2**).

**Table 2.**
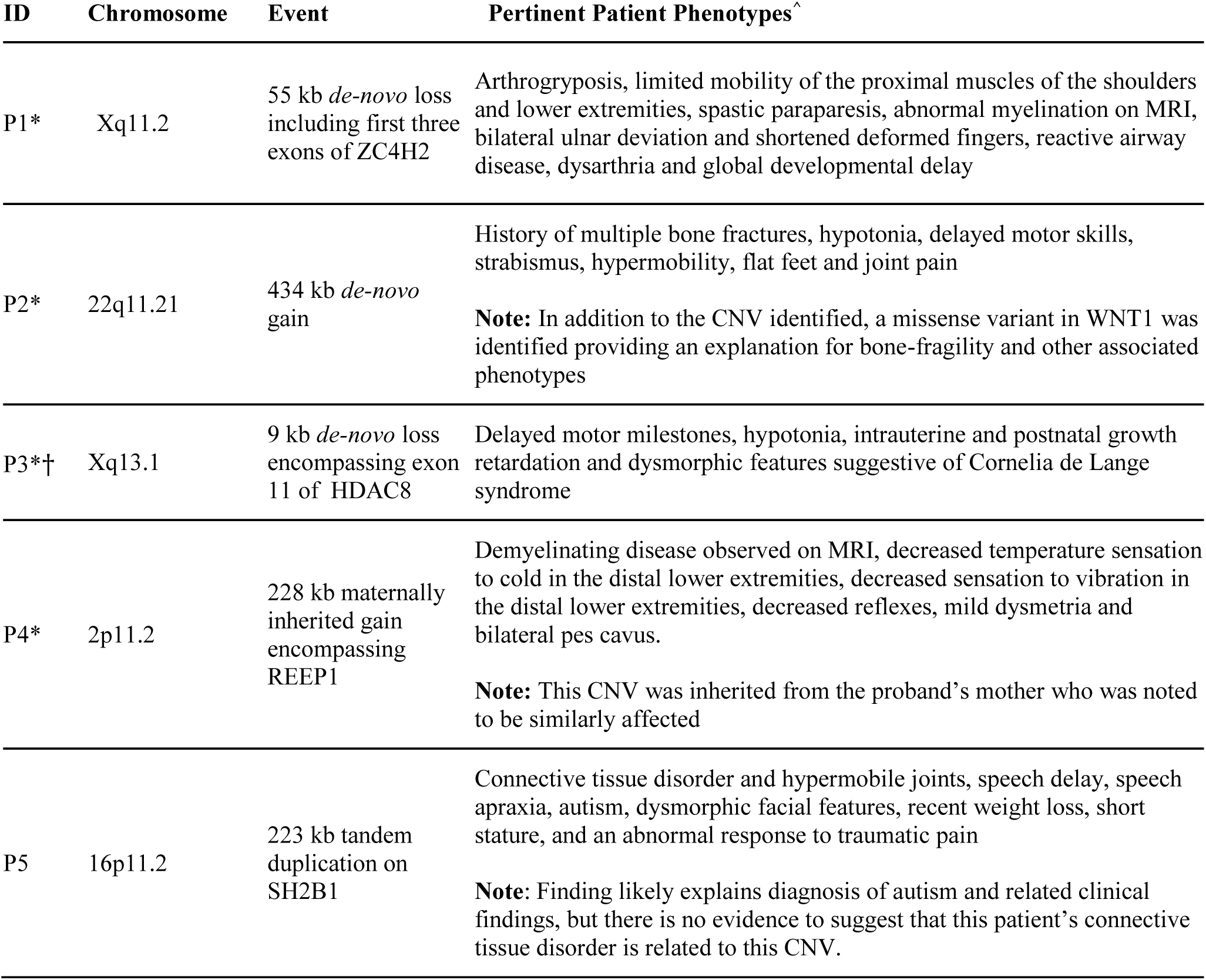
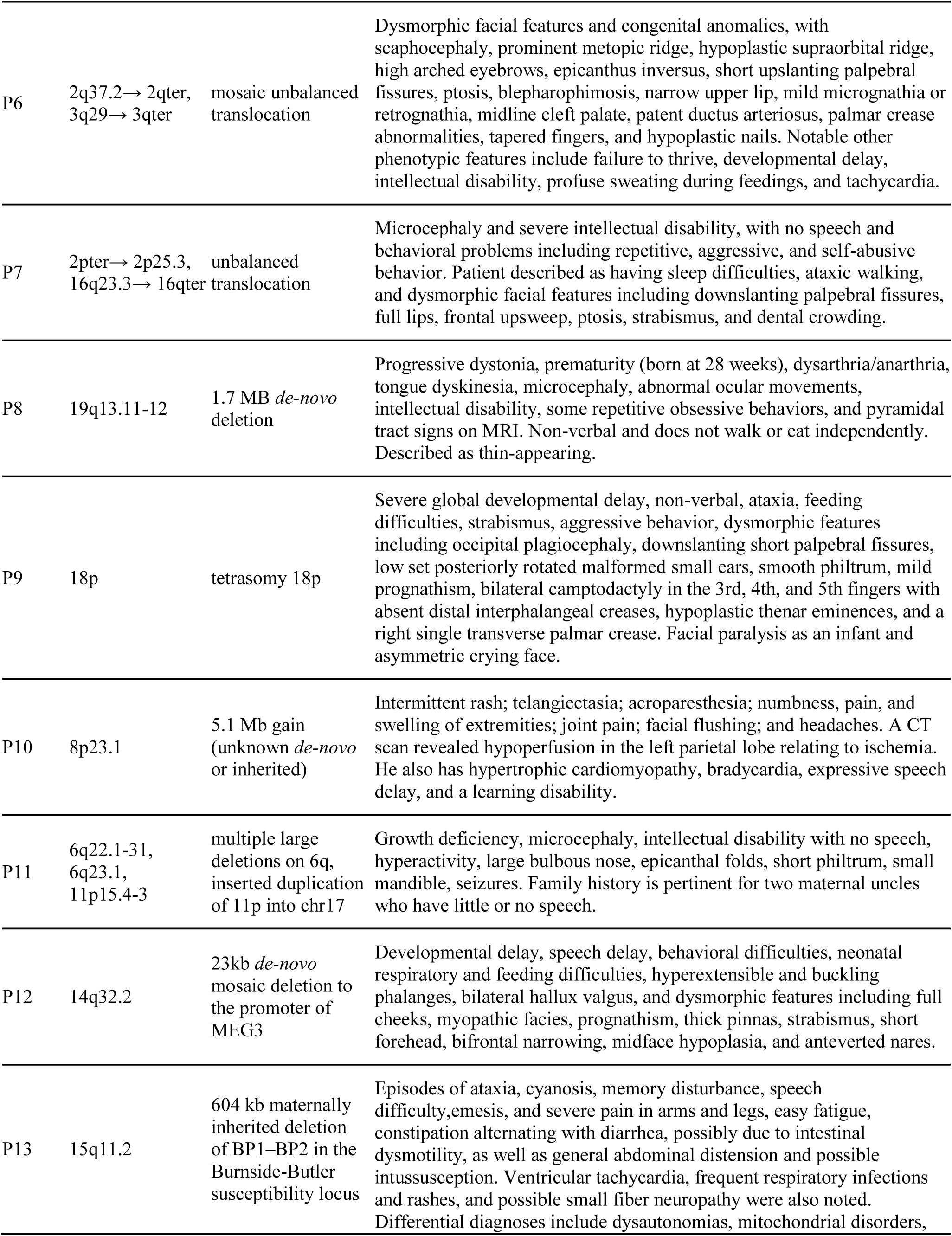
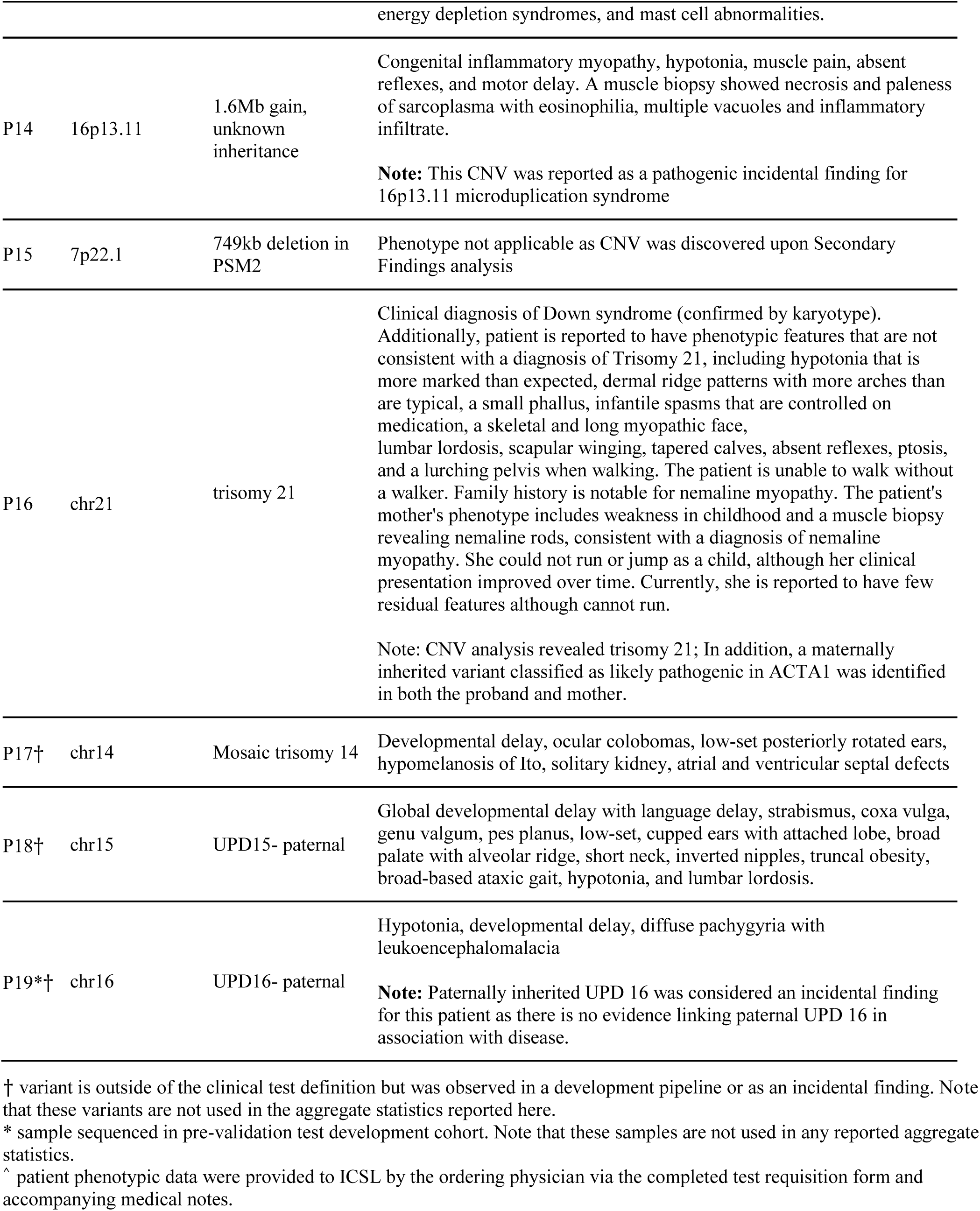
Summary of clinically relevant CNVs.

### Resolution of complex CNVs

We found that the combination of depth-based CNV calling and the utilization of discordant read-pair information and reassembly of breakpoints can enable deconvolution of complex rearrangements. In one example, family based CNV analysis of case P1 identified a 55kb *de-novo* deletion of the first three exons of *ZC4H2* on the paternal X chromosome (**Figure 1**), consistent with Wieacker-Wolff syndrome (**Table 2**). Structural variant analysis (**Methods**) identified evidence for a tandem duplication in the proband’s father, sharing a breakpoint with the deletion in the proband (**Figure 1b**). Read-depth information from the father shows a copy-number gain directly upstream of the *de-novo* deletion in the proband (**Figure 1a**). Taken together these data likely indicate a multi-stage repair mechanism contributing to the copy-number loss in the proband.

**Figure 1.**
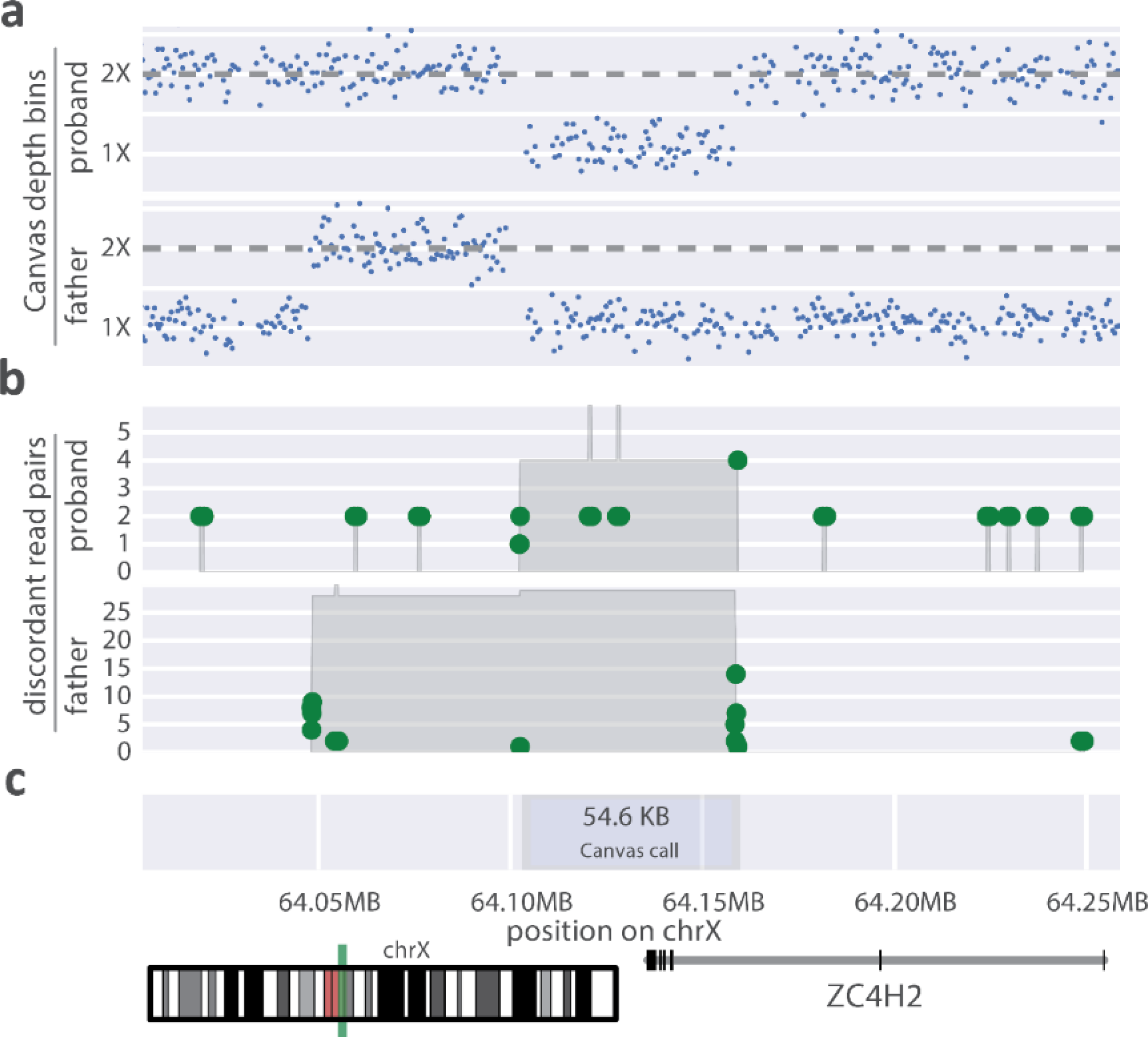
ZC4H2 *de-novo* deletion in case **P1**. a) Normalized sequencing depth for proband and her father. b) Location of discordant read pairs (>1000bp insert size), where green dots represent the location of paired ends in a discordant read-pair, the grey shaded area represents the total number of discordant read-pairs spanning a given genomic segments. c) Annotation of the original Canvas call boundaries as well as the location of the CNV on chromosome X.

Similarly, case P11 harbored multiple CNVs indicative of a large chromosomal disruption event (Fukami et al., 2017; Liu et al., 2017) including 15.5 MB and 2.5MB deletions on 6q along with a 2MB copy-number gain on 11p (**Figure S4a, Table 2**). Inspection of structural variation near these events yielded discordant reads spanning the two deletions, supporting the presence of simple deletions as opposed to more complex events such as translocations or inversions (**Figure S4b**), however such complex structural variation cannot be definitively ruled out. Structural variation near the boundaries of the 11p gain indicated an insertion of this duplicated DNA segment into 17q21.3 (**Figure S4c**).

Copy number analysis of case P7 (**Table 2**) identified a 7MB *de-novo* terminal duplication on 16q and a 3MB *de-novo* terminal deletion on 2p (**Figure 2a-b**). Analysis of variant allele frequencies overlapping these two CNVs phased both alterations to the paternal chromosome (**Figure 2c-d, Methods, Supplemental Note**). Subsequent analysis of sequencing reads provided evidence for a balanced translocation in the father as well as the unaffected sister, while the proband had support for an unbalanced translocation (**Figure 2e, Figure S5, Methods**).

**Figure 2.**
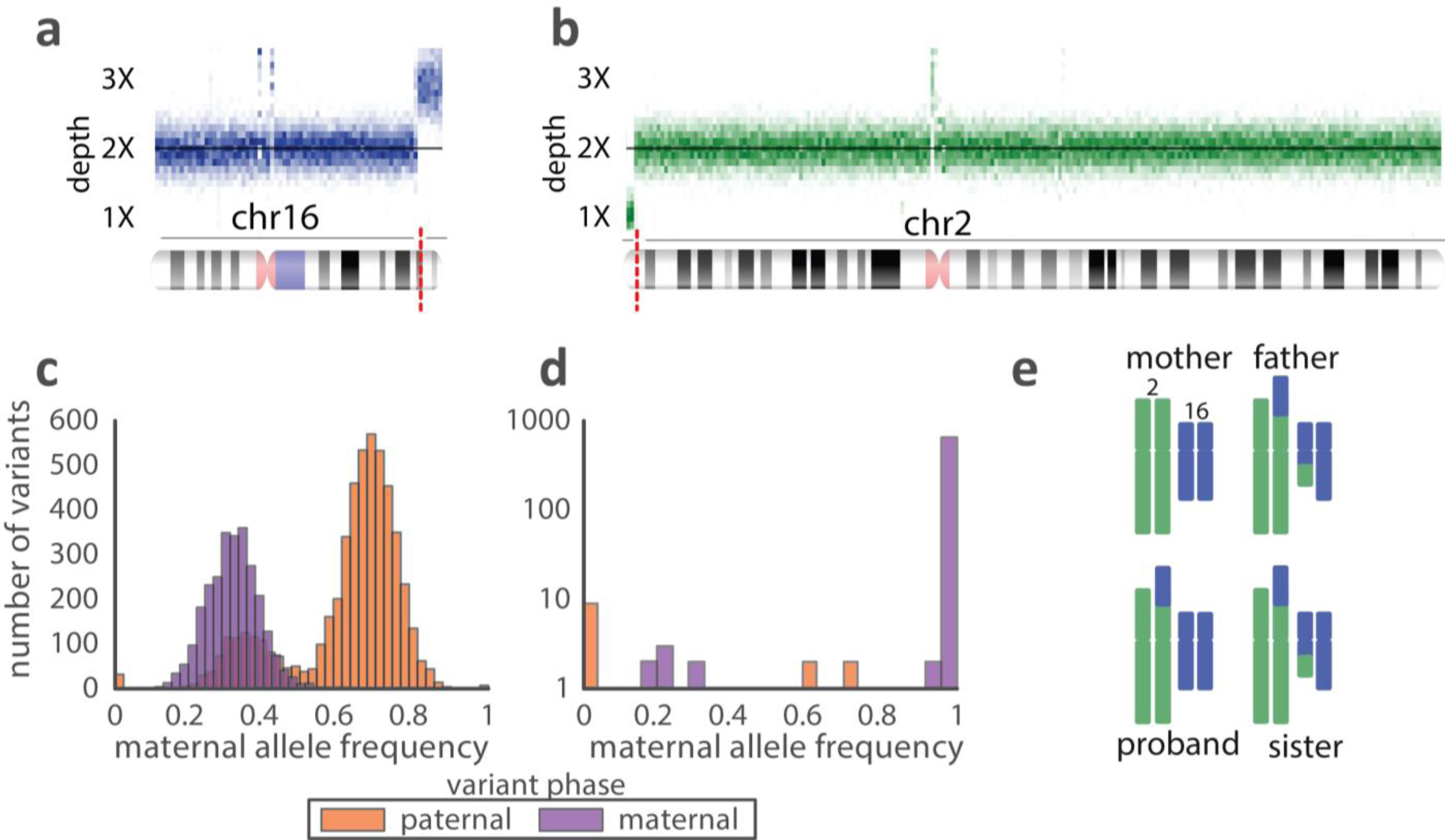
Case P7 - derivative chromosome inherited from a balanced translocation in a parent. **a-b**) Sequencing depth support for duplication on 16q (**a**) and deletion on 2p (**b**). Slices in the image represent distribution of normalized sequencing depth across 100kb genomic intervals. **c-d**) Distribution of maternal allele frequency for all phased variants in copy-number altered regions corresponding to 16q gain (**c**) and 2p loss (**d**). Note that variant frequency distributions are colored by the parent of origin as determined by trio phasing. **e**) Summary of split and discordant sequencing read evidence for recombinant chromosomes at CNV breakpoints.

**Figure 3.**
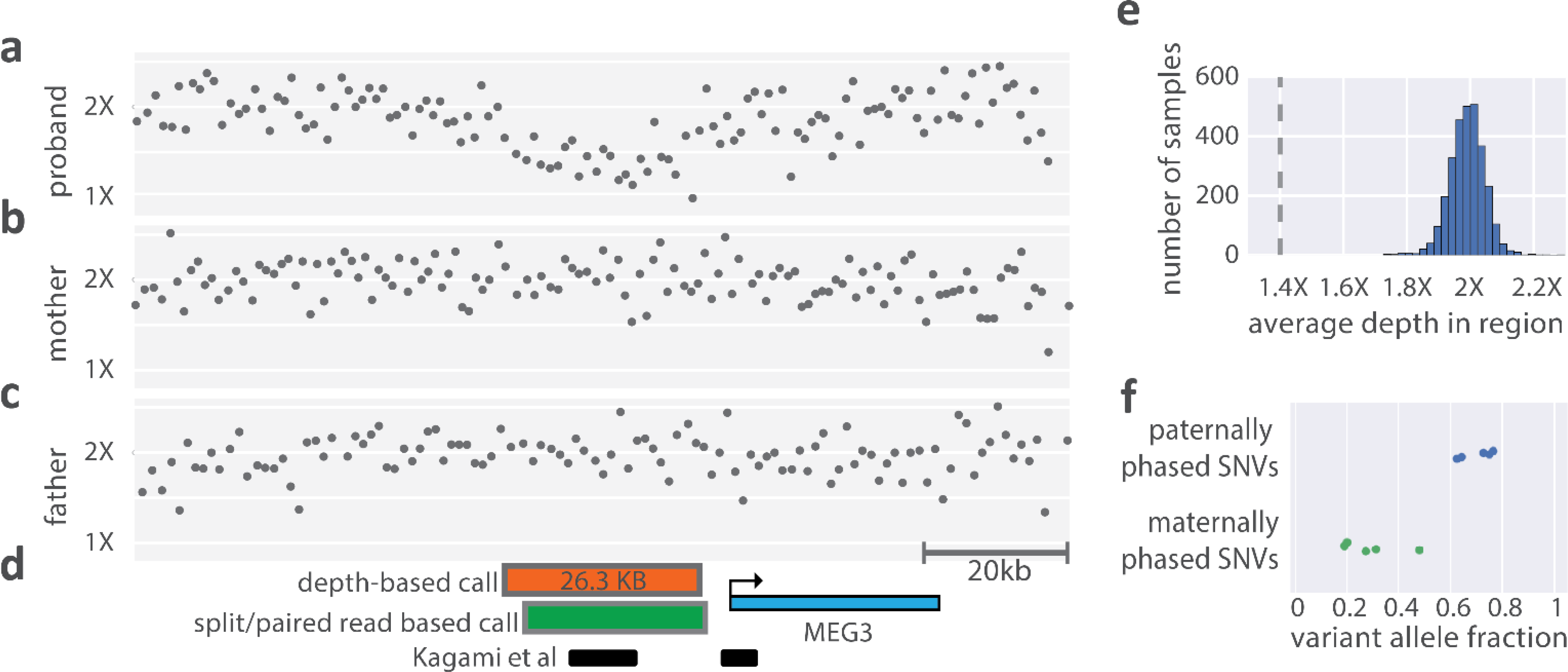
Mosaic 14q32.2 26kb microdeletion. **a-c**, Normalized depth across pedigree sequenced for subject P12. Shown here is the genomic region between 101.21MB and 101.34MB on chromosome 14 (hg19 coordinates) **d**, Annotations for the genomic region. The orange box represents the Canvas CNV call boundaries, the green box represents breakpoint assembled coordinates of the deletion from the Manta structural variant (SV) caller, the black lines represent subjects from Kagami et al (Kagami et al., 2010) with deletions in this region, the blue box represents the gene boundaries of the imprinted gene MEG3. **e**, Average depth across the region of the CNV call for samples across an internal reference population, depth for the proband is indicated with a horizontal dashed line. **f**, Variant allele frequency for the SNVs within the deleted region.

In case P6 we observed a similar unbalanced translocation with a non-homologous break-end linking the centromeric breakpoints of the two large terminal CNVs. Further inspection of copy-number depth as well as variant allele frequencies indicated that the CNVs were likely mosaic in the blood, with both events having similar estimated purity of 60-64% purity of the sample (**Figure S6**), which external testing confirmed at 63%. Taken together these events suggest the presence of a mosaic unbalanced translocation in the affected proband, a rare event which has been proposed to occur by various mechanisms of recombination (Gijsbers et al., 2011).

### cWGS identifies a clinically relevant mosaic non-coding CNV

In case P12, standard CNV analysis identified a 23kb *de-novo* deletion upstream of MEG3 completely overlapping the IG-DMR region of 14q32.2 previously implicated in Kagami-Ogata syndrome (Kagami et al., 2010). Further analysis indicated that the CNV was likely mosaic, present in about 50% of cells, and that the deletion phased to the maternal allele, consistent with the paternal imprinting mechanism of Kagami-Ogata (Kagami et al., 2005). In this case, the mosaic deletion passed standard CNV analysis and was annotated as mosaic during manual curation. In contrast, in case P17, we identified a mosaic trisomy of chromosome 14 via a genome-wide visualization (**Figure S7**). While this mosaic variant was identified outside of our clinically validated pipeline, the variant was sent out for external testing which confirmed and estimated its purity at 51%, compared to 47% as estimated by WGS.

## Discussion

Here we report the development, validation, and deployment of a multifaceted clinical test for individuals with rare and undiagnosed disease (RUGD). With whole genome sequencing, copy number variation can be profiled alongside small variant analysis without any additional sample preparation or experimental protocols. Due to the purely bioinformatic nature of this addition, we have been able to synchronize analysis and reporting of these multiple classes of variants.

In our CNV calling pipeline, we optimize the parameters of the caller to favor sensitivity (**Methods**). In our validation, this provided greater recovery of externally annotated CNVs than clinical microarrays (**Table 1**), but may also result in an increased false-positive rate. To address this, we have developed a stringent filtering and manual curation protocol (**Methods**, **Figure S1-3**). This curation relies heavily on our ability to annotate population frequency (**Figure S2d**), as well as visualization of the CNV call to assess the underlying data quality (**Figure S3**). In addition, we leverage external databases of benign and pathogenic CNVs (Firth et al., 2009; MacDonald et al., 2014), internal aggregate data, and previously curated variants to assess the analytical validity of a call and provide a variant classification (Kearney et al., 2011).

These methods do not rely on bulk data processing or analysis (i.e., batching of samples), allowing for ingestion and interpretation of one family at a time. This addresses the time-lag from sample collection to interpretation present in some laboratory-based clinical CNV analysis protocols. Furthermore, these methods are suited to exploit future increases in sequencing coverage that will result in an increase in resolution to call small CNVs, allowing for test improvement with minimal modifications to the sample-preparation or bioinformatic pipelines.

In many families with previous genetic testing, cWGS was able to identify new variants and provide a diagnosis. For example, in the case of a child from a resource limited clinic in South America (P14, **Table 2**) who had a previous negative clinical exome, cWGS was able to identify a pathogenic 1.7MB deletion indicative of 16p13.11 Microdeletion Syndrome. In subject P16 (**Table 2**) we observed trisomy 21 in the subject consistent with a pre-existing Down Syndrome diagnosis, but were also able to identify a likely pathogenic SNV within the ACTA1 gene conferring the additional diagnosis of an inherited nemaline myopathy.

In a select number of cases, the cWGS CNV pipeline was able to identify variants that would have been missed by exome, single gene testing and microarray. Most notable of these was the deletion in P12, which is below the limit of most commercial microarrays (26kb), sits in a non-coding region upstream of a long non-coding RNA (MEG3), occurs in a locus that is paternally imprinted, phases to the maternal chromosome and is 50% mosaic. We are not aware of another agnostic genome-wide testing approach which would have been able to identify this variant and capture all the associated features.

Future test improvements will focus on bringing a broader diversity of variants within the umbrella of cWGS. Identification of mosaic CNVs remains a priority, especially for large copy-number events such as trisomy where cWGS has sufficient data to detect low purity alterations (Dong et al., 2016). Structural variant calling will be needed to fill the currently existing gap between small variants (SNVs and INDELs) and depth-based CNVs. Uniparental isodisomy and heterodisomy, which we have previously seen as incidental findings (**Table 2**, case P18 and P19), will be incorporated via observation of inheritance patterns of small variants. Finally, validation of specialized variant callers to detect hard-to-call variation such as repeat expansions (Dolzhenko et al., 2016; Gatchel and Zoghbi, 2005) and SMA (Feng et al., 2017) from cWGS data will open up the test to new classes of disease.

In conclusion, we present our experience of the development and deployment of CNV calling on top of an existing cWGS assay. As sequencing costs continue to decrease, the use of a first line whole genome diagnostic spanning a broad spectrum of genetic variation will become the standard for rare and undiagnosed disease.

## Acknowledgments

We would like to thank the families who participated in this study. Much of the sequencing in this study was made possible by the Illumina foundation as part of the iHope program, which donates clinical WGS to underserved families with rare and undiagnosed disease. We would also like to thank Dr. Marilyn Jones, Ms. Diane Masser-Frye, Dr. Adeline Vanderver, along with our clinical collaborators, as well as the work of the Illumina Clinical Services Laboratory in sample processing, sequencing and data management.

## Author contributions

**Case management:** Carolyn Brown, Krista Bluske, Nicole Burns, Aditi Chawla, Alison J Coffey, Alka Malhotra, Alicia Scocchia, Erin Thorpe, R Tanner Hagelstrom, Denise L Perry

**Bioinformatic analysis:** Andrew M Gross, Vani Rajan, Subramanian S. Ajay, Egor Dolzhenko, Bryan Lajoie, Michael Eberle

**Clinical microarray analysis:** Natasa Dzidic, Karine Hovanes, Trilochan Sahoo

**Aggregate data analysis:** Andrew M Gross

**Study design/Manuscript writing:** Andrew M Gross, Denise L Perry, Subramanian S. Ajay, R Tanner Hagelstrom, John Belmont, Ryan J Taft

**Study coordination:** Julia McEachern

## Supplemental Figures

**Figure S1.**
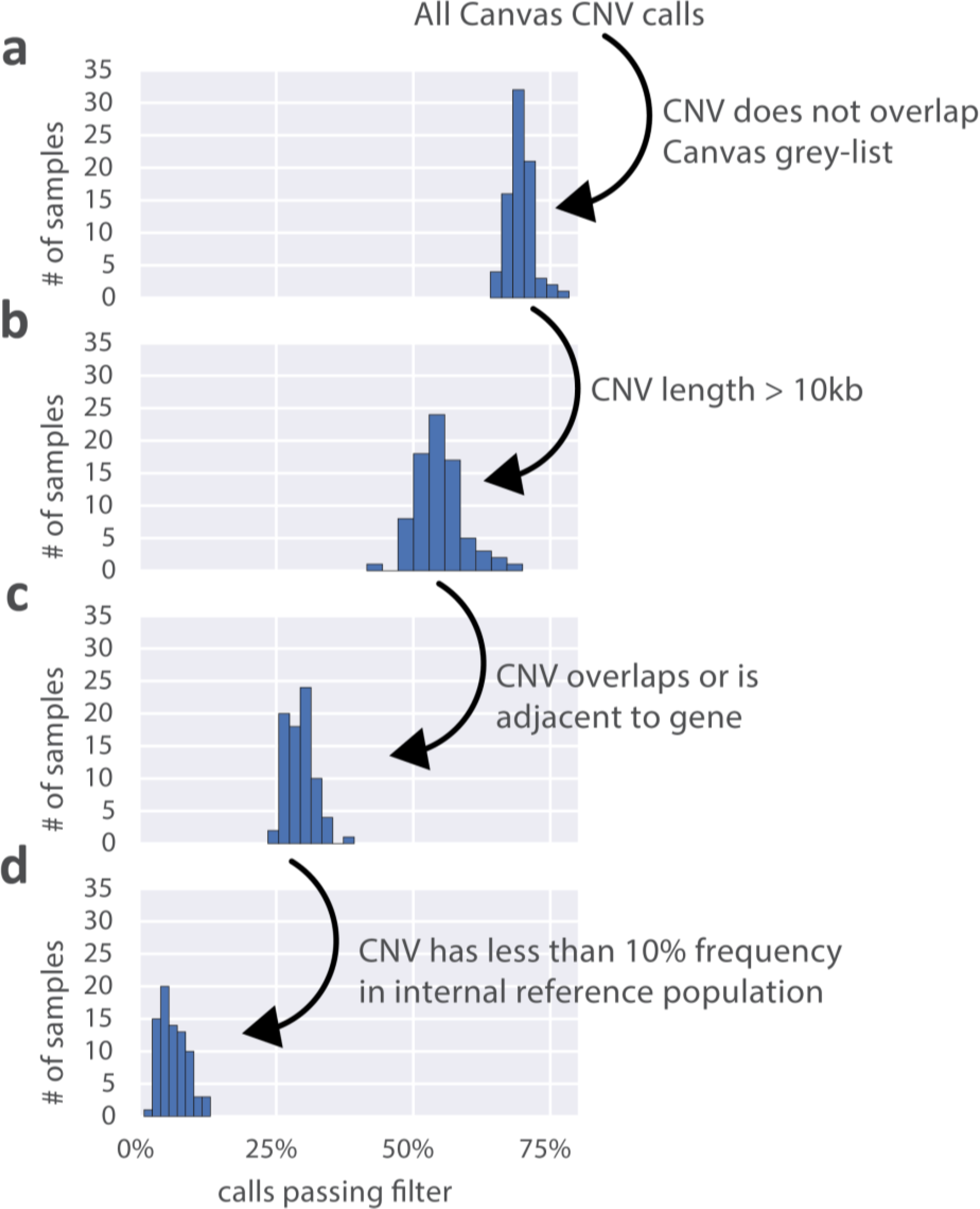
CNV filtering pipeline. During data processing, automated filters are applied to call-sets to limit the number of calls presented to case managers for manual curation. Shown in each panel are the percentage of calls remaining after each filtering step is applied sequentially. Distributions reflect 79 samples assessed for CNVs in the ICSL cohort. For details on individual filters, see **Methods**.

**Figure S2.**
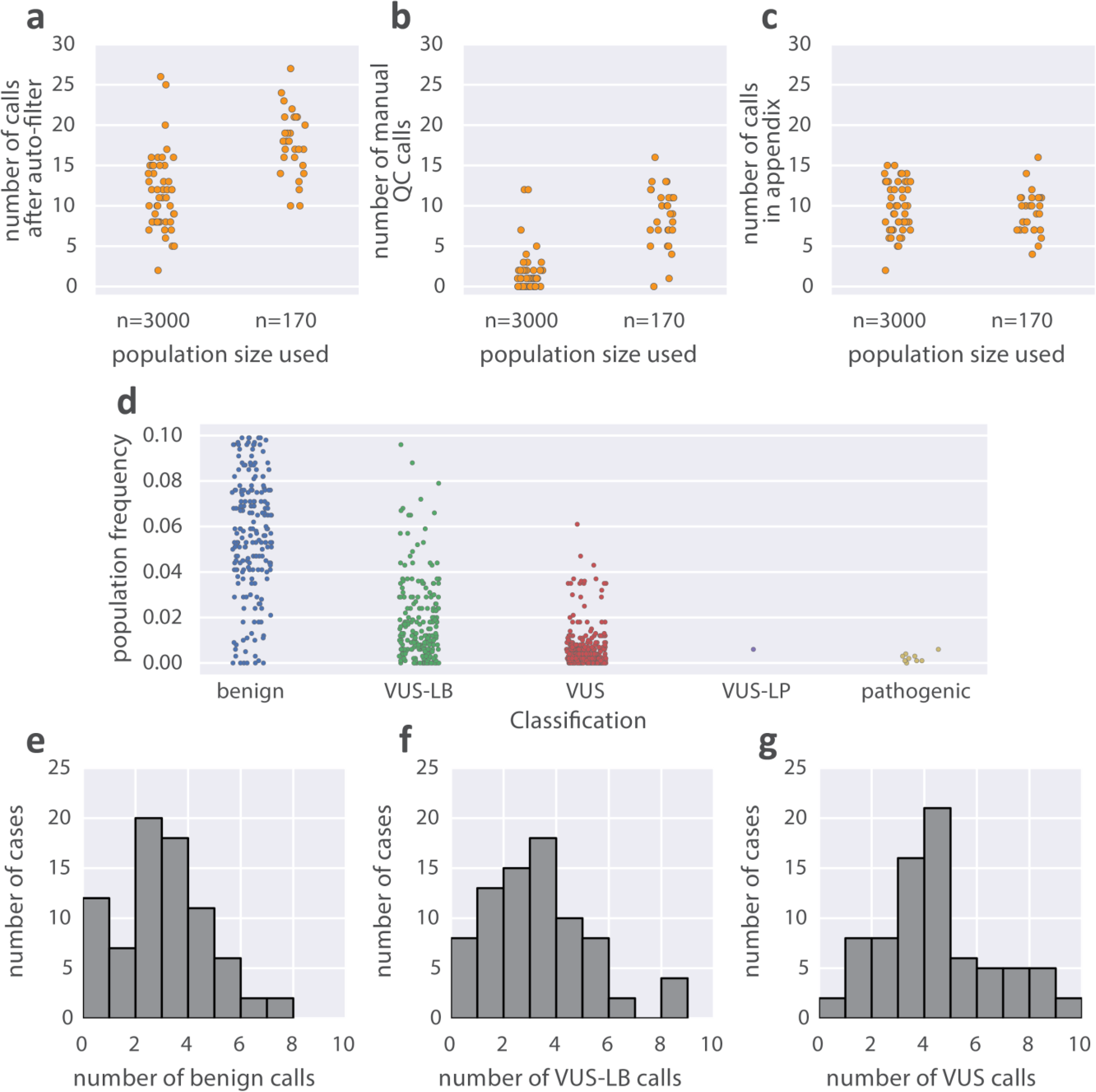
Summary of manual curation and variant annotation across the CNV clinical case cohort. (**a-c**) Number of calls passing automated filters (**a**), manually filtered (**b**) and included in a CNV appendix to clinical reports (**c**) broken down by the two populations used for CNV frequency annotation (n=170 was used for the first 28, and n=3000 was used for the remainder). (**d**) Copy number variant population frequency broken down by variant classification post curation. (**e-g**) Breakdown of variant classifications for CNVs included in clinical report appendix across the cohort. In addition there were calls reported as pathogenic in 7 cases, and one likely-pathogenic call reported. VUS - variant of unknown significance; VUS-LB-variant of unknown significance, likely benign.

**Figure S3.**
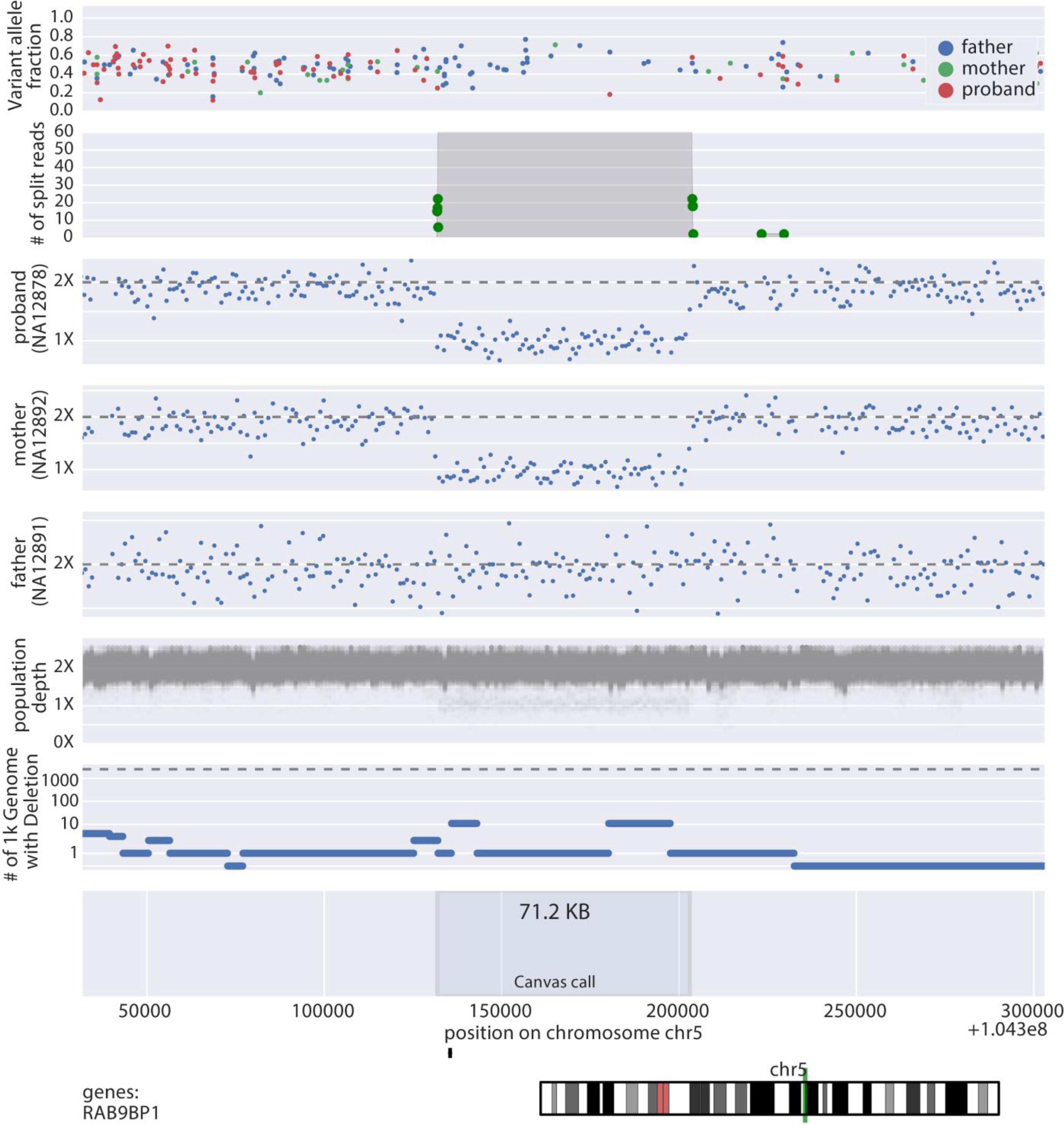
Example CNV visualization used for CNV interpretation. **Variant allele fraction**: read-count ratio of two alleles for all heterozygous SNVs in a region. This fraction should be centered at 1/2 in diploid regions, while for duplications it is expected to be centered around 1/3 and 2/3 as there is an imbalance of alleles. For deletions there should be an absence of heterozygous SNVs due to the presence of only one allele. **Number of Split Reads:** Green dots show the locations of discordant read-pairs, while the height of the grey shaded region indicates the number of discordant reads spanning across a given region on the genome. While this is a useful confirmation for CNVs breakpoints in many cases, CNVs may have breakpoints in non-unique sequence resulting in an inability to uniquely map reads to the flanks of the CNV. **Read depth:** Normalized read depth across the proband and parents. This allows for evaluation of error modes in the caller and may expose patterns not yet be picked up by an automated caller such as inheritance of a mosaic CNV. **1000 Genomes Data:** CNV calls the 1000 Genomes Project (WGS, 7x coverage) (Sudmant et al., 2015) used to identify common deletions. **Population Depth Data:** Normalized coverage for 200 samples selected from our internal population data. This allows for inspection of population trends which may expose artifacts in the read-mapping or data-normalization process leading to a false-positive call. **Chromosome view and overlapping genes:** This field allows the interpreter to view where the event takes place in context of the chromosome and displays the coding sequences of genes that overlap the events so it can be determined if the genes are relevant to the phenotype.

**Figure S4.**
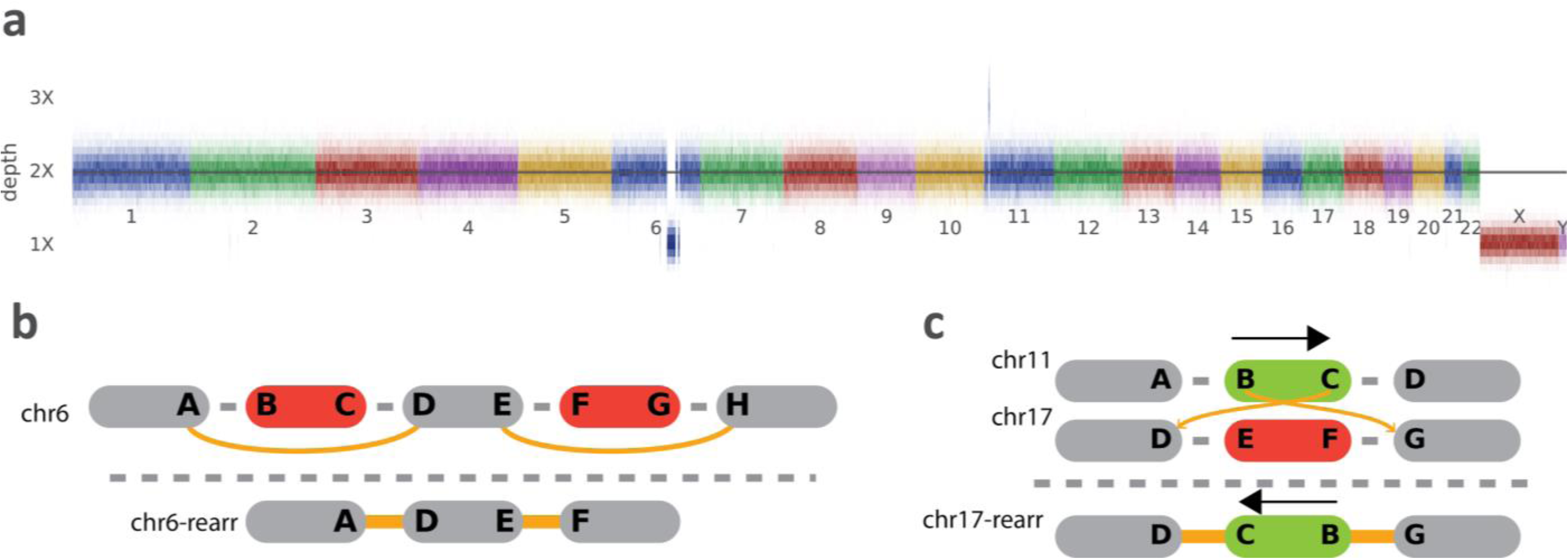
Complex rearrangement in subject P11. **a**, Depth across chromosomes showing two large regions of copy-number loss on chromosome 6 and a copy number gain on chromosome 11. **b**, Schematic of structural rearrangement on chromosome 6 indicating two large deletions in close proximity. **c**, Schematic of structural rearrangement on chromosome 17 indicated a large insertion of genetic material from chromosome 11.

**Figure S5.**
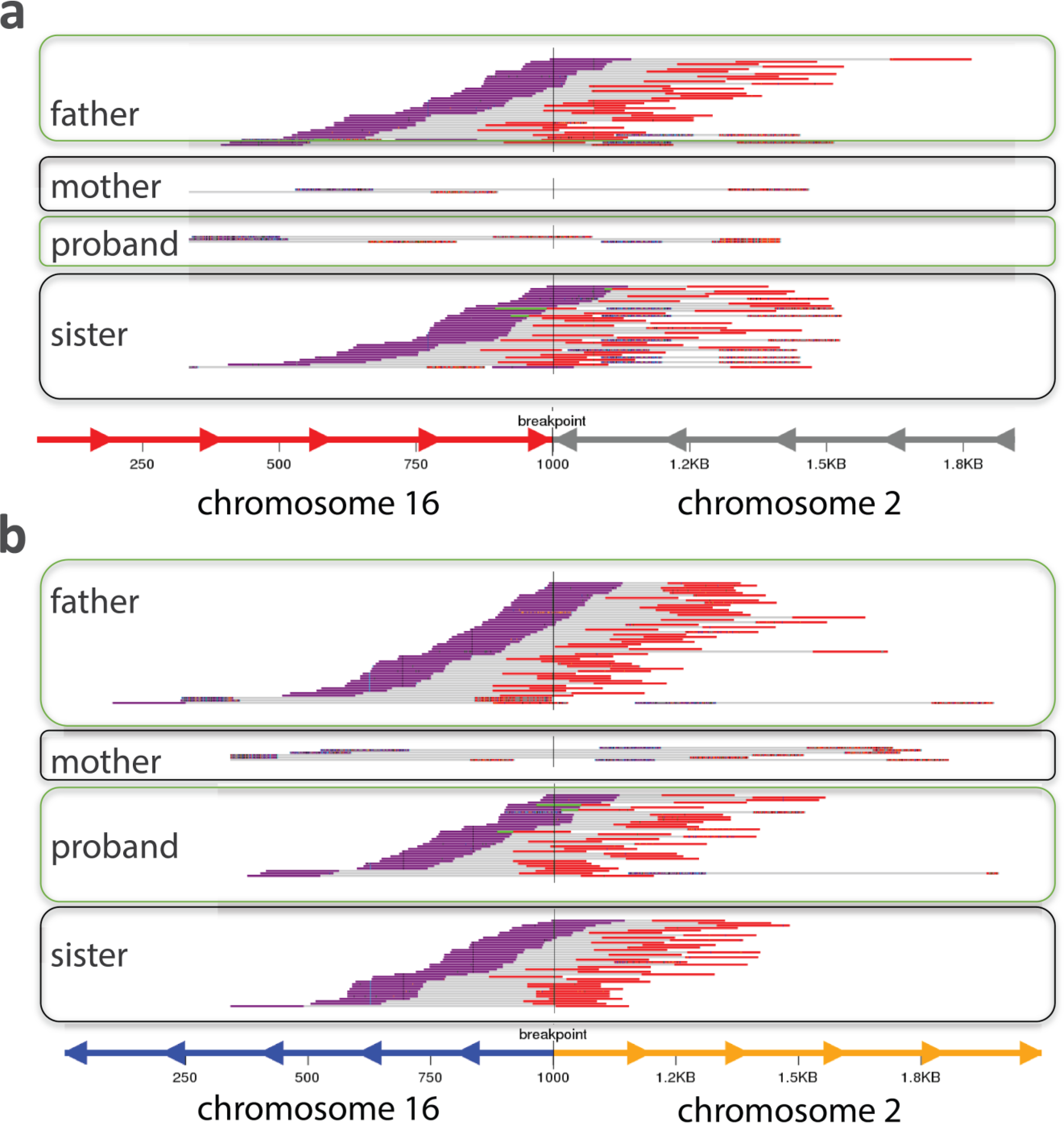
Read support for unbalanced translocation in subject P7. Shown here are modified plots from the svviz graph realignment program. In brief reads are realigned to normal (not shown) and recombinant (shown here) chromosomes across the pedigree. Purple and red colors represent the first and second reads in the read-pair, respectively, for details on the visualization and realignment methods see (Spies et al., 2015). See also **Figure 2**.

**Figure S6.**
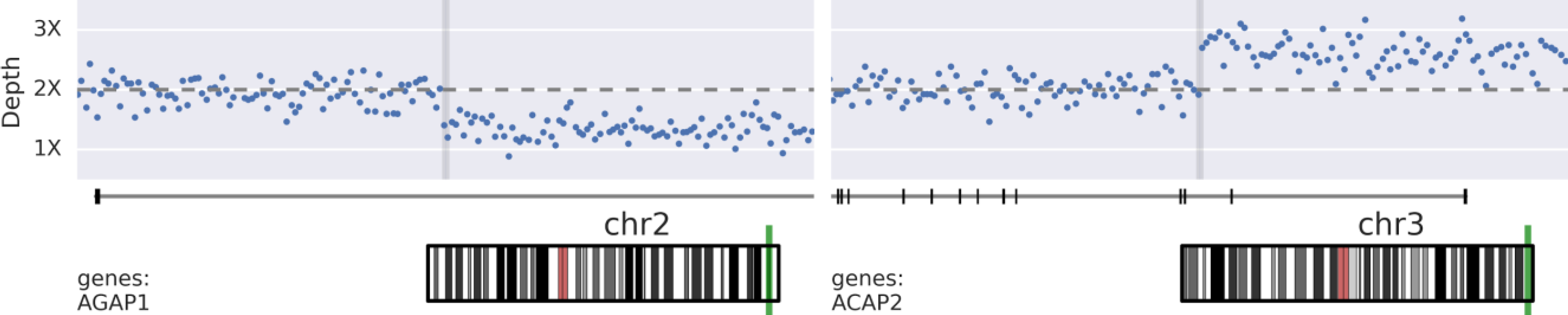
Depth at CNV breakpoints for a mosaic unbalanced translocation in subject P6. Horizontal grey lines correspond to the location of a non-homologous chromosomal break-end uncovered in structural variant analysis.

**Figure S7.**
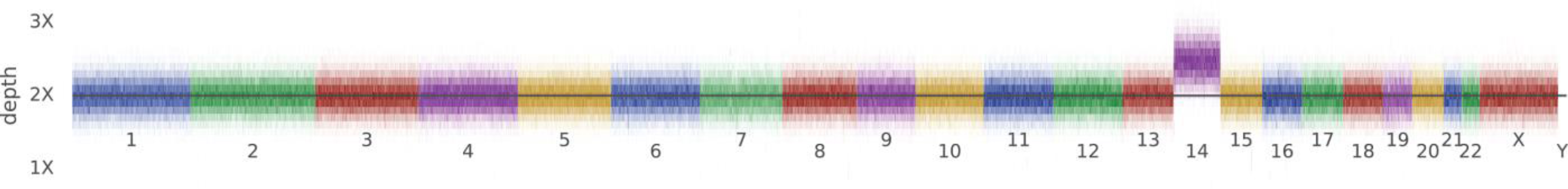
Depth across chromosomes for subject P6 with mosaic trisomy 14. Horizontal line corresponds to diploid copy-number.

**Table S1:**
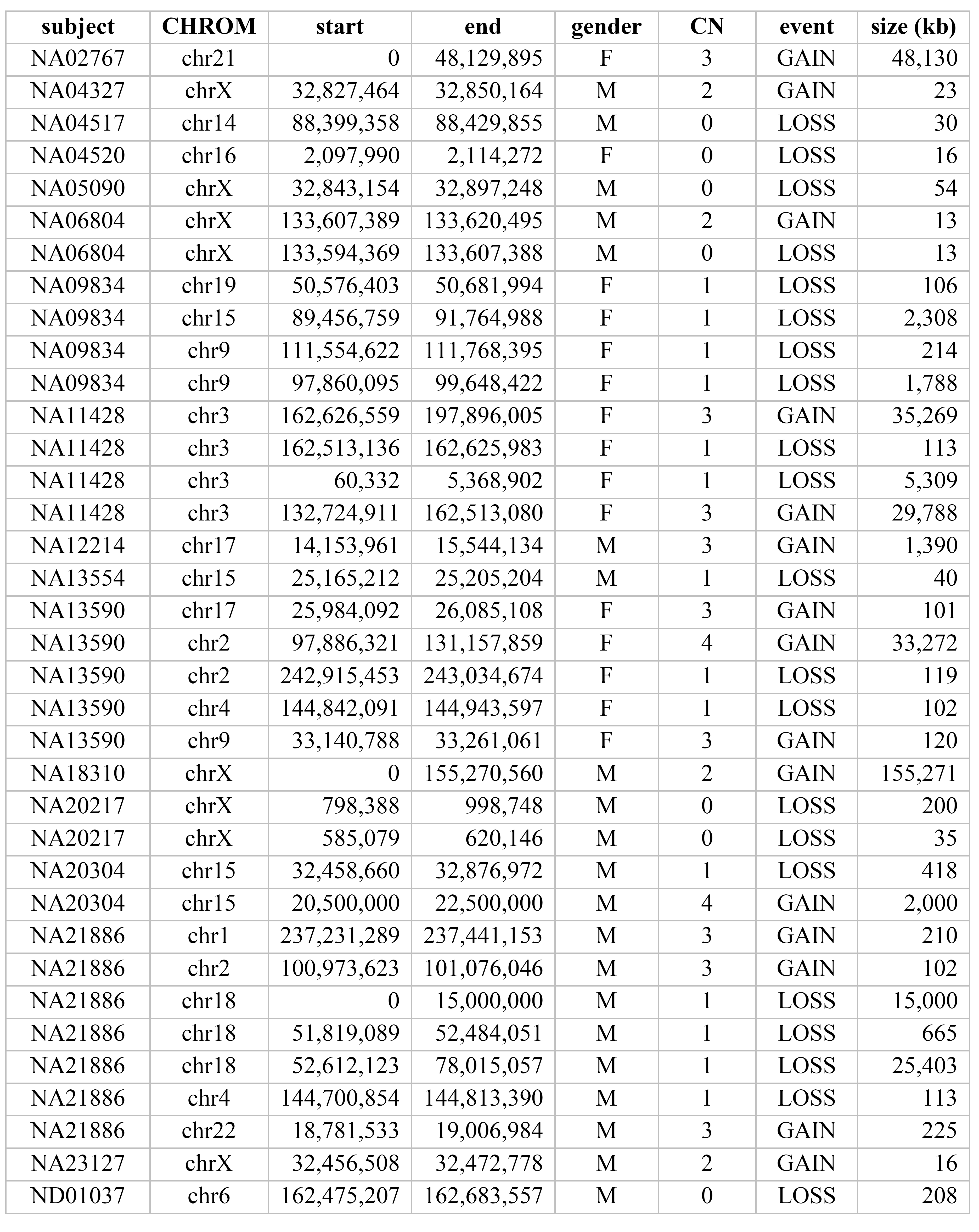
Coriell reference CNV calls.

| subject | CHROM | start | end | gender | CN | event | size (kb) |
| --- | --- | --- | --- | --- | --- | --- | --- |
| NA02767 | chr21 | 0 | 48,129,895 | F | 3 | GAIN | 48,130 |
| NA04327 | chrX | 32,827,464 | 32,850,164 | M | 2 | GAIN | 23 |
| NA04517 | chr14 | 88,399,358 | 88,429,855 | M | 0 | LOSS | 30 |
| NA04520 | chr16 | 2,097,990 | 2,114,272 | F | 0 | LOSS | 16 |
| NA05090 | chrX | 32,843,154 | 32,897,248 | M | 0 | LOSS | 54 |
| NA06804 | chrX | 133,607,389 | 133,620,495 | M | 2 | GAIN | 13 |
| NA06804 | chrX | 133,594,369 | 133,607,388 | M | 0 | LOSS | 13 |
| NA09834 | chr19 | 50,576,403 | 50,681,994 | F | 1 | LOSS | 106 |
| NA09834 | chr15 | 89,456,759 | 91,764,988 | F | 1 | LOSS | 2,308 |
| NA09834 | chr9 | 111,554,622 | 111,768,395 | F | 1 | LOSS | 214 |
| NA09834 | chr9 | 97,860,095 | 99,648,422 | F | 1 | LOSS | 1,788 |
| NA11428 | chr3 | 162,626,559 | 197,896,005 | F | 3 | GAIN | 35,269 |
| NA11428 | chr3 | 162,513,136 | 162,625,983 | F | 1 | LOSS | 113 |
| NA11428 | chr3 | 60,332 | 5,368,902 | F | 1 | LOSS | 5,309 |
| NA11428 | chr3 | 132,724,911 | 162,513,080 | F | 3 | GAIN | 29,788 |
| NA12214 | chr17 | 14,153,961 | 15,544,134 | M | 3 | GAIN | 1,390 |
| NA13554 | chr15 | 25,165,212 | 25,205,204 | M | 1 | LOSS | 40 |
| NA13590 | chr17 | 25,984,092 | 26,085,108 | F | 3 | GAIN | 101 |
| NA13590 | chr2 | 97,886,321 | 131,157,859 | F | 4 | GAIN | 33,272 |
| NA13590 | chr2 | 242,915,453 | 243,034,674 | F | 1 | LOSS | 119 |
| NA13590 | chr4 | 144,842,091 | 144,943,597 | F | 1 | LOSS | 102 |
| NA13590 | chr9 | 33,140,788 | 33,261,061 | F | 3 | GAIN | 120 |
| NA18310 | chrX | 0 | 155,270,560 | M | 2 | GAIN | 155,271 |
| NA20217 | chrX | 798,388 | 998,748 | M | 0 | LOSS | 200 |
| NA20217 | chrX | 585,079 | 620,146 | M | 0 | LOSS | 35 |
| NA20304 | chr15 | 32,458,660 | 32,876,972 | M | 1 | LOSS | 418 |
| NA20304 | chr15 | 20,500,000 | 22,500,000 | M | 4 | GAIN | 2,000 |
| NA21886 | chr1 | 237,231,289 | 237,441,153 | M | 3 | GAIN | 210 |
| NA21886 | chr2 | 100,973,623 | 101,076,046 | M | 3 | GAIN | 102 |
| NA21886 | chr18 | 0 | 15,000,000 | M | 1 | LOSS | 15,000 |
| NA21886 | chr18 | 51,819,089 | 52,484,051 | M | 1 | LOSS | 665 |
| NA21886 | chr18 | 52,612,123 | 78,015,057 | M | 1 | LOSS | 25,403 |
| NA21886 | chr4 | 144,700,854 | 144,813,390 | M | 1 | LOSS | 113 |
| NA21886 | chr22 | 18,781,533 | 19,006,984 | M | 3 | GAIN | 225 |
| NA23127 | chrX | 32,456,508 | 32,472,778 | M | 2 | GAIN | 16 |
| ND01037 | chr6 | 162,475,207 | 162,683,557 | M | 0 | LOSS | 208 |

**Table S2.** Summary of sensitivity of cWGS and clinical microarrays to annotated CNVs in cell-lines.

|  |  | Coriell Events | Called by<br>Array**<br>(curated) | Called by<br>WGS** |
| --- | --- | --- | --- | --- |
| event | size |  |  |  |
| LOSS | 10kb-50kb | 5 | 3 (+1) | 4 |
|  | 50kb-100kb | 1 | 1 | 1 |
|  | 100kb-500kb | 9 | 2 (+1) | 5 |
|  | >500kb | 6 | 6 | 6 |
|  | Overall | 21 | 12 (57%) | 16 (76%) |
| GAIN | 10kb-50kb | 3 | 0 | 2 |
|  | 100kb-500kb | 5 | 4 | 4 |
|  | >500kb | 7 | 4 | 5 |
|  | Overall | 15 | 8 (53%) | 11 (73%) |
| All CNV calls |  | n=36 | 20 (56%) | 27 (75%) |
| **75% overlap<br>Note that +1 indicates calls that were not in call set, but recovered in manual review |  |  |  |  |

**Table S3.** Coordinates of reported variants from RUGD cases.

| ID | event | CHROM | CNV coordinates |  | SV coordinates |  | comment |
| --- | --- | --- | --- | --- | --- | --- | --- |
|  |  |  | start | end | start | end |  |
| P1 | LOSS | chrX | 64104162 | 64158754 |  |  | split read evidence but no SV call |
| P2 | GAIN | chr22 | 21052009 | 21484438 |  |  |  |
| P3 | LOSS | chrX | 71549289 | 71557651 | 71549289 | 71557651 | CNV came from development SV pipeline |
| P4 | GAIN | chr2 | 86283023 | 86511034 | 86282714 | 86510931 |  |
| P5 | GAIN | chr16 | 28,823,31 | 29047087 |  |  |  |
| P6 | LOSS | chr2 | 236478472 | 243048854 | 236478812 |  | breakpoint links CNVs in unbalanced translocation |
| P6 | GAIN | chr3 | 195106447 | 197846145 | 195105935 |  | breakpoint links CNVs in unbalanced translocation |
| P7 | LOSS | chr2 | 11314 | 3033976 |  | 3033857 | breakpoint links CNVs in unbalanced translocation |
| P7 | GAIN | chr16 | 82865402 | 90163542 | 82865480 |  | breakpoint links CNVs in unbalanced translocation |
| P8 | LOSS | chr19 | 35223021 | 36895699 | 35223614 | 36896374 |  |
| P9 | GAIN | chr18 | 11494 | 15404287 |  |  |  |
| P10 | GAIN | chr8 | 7153587 | 12245784 |  |  |  |
| P11 | LOSS | chr6 | 109324789 | 124836619 | 109325818 | 124836270 |  |
| P11 | LOSS | chr6 | 129969121 | 132499298 | 129970203 | 132499992 |  |
| P11 | GAIN | chr11 | 8548056 | 10497905 | 8548078 | 10498608 | Inserted into chr17 |
| P11 | INSERTION | chr17 |  |  | 41705963 | 41705972 | Genomic material from chr11 inserted here |
| P12 | LOSS | chr14 | 101261679 | 101288013 | 101261679 | 101288013 |  |
| P13 | LOSS | chr15 | 22696624 | 23301066 |  |  |  |
| P14 | LOSS | chr16 | 14800000 | 16400000 |  |  | CNV boundaries approximate due to low sequence complexity on flanks |
| P15 | LOSS | chr7 | 6027017 | 6776186 |  |  |  |
| P16 | GAIN | chr21 | 14596056 | 48101324 |  |  |  |
| P17 | GAIN | chr14 |  |  |  |  | CNV identified by visualization and boundaries manually assessed |
| P18 | ROH | chr15 | 23633319 | 102280298 |  |  | boundaries assessed by ROH caller |
| P19 | ROH | chr16 | 80212 | 90142842 |  |  | boundaries assessed by ROH caller |

## Supplemental Note: Manual Inspection of NA12878 calls with partial BioNano overlap

In these cases, calls from our WGS call-set had partial overlap with the BioNano calls. We inspected these manually to better understand the discrepancies and assess false-positive or true-positive status.

**Supplemental Note Figure 1.**
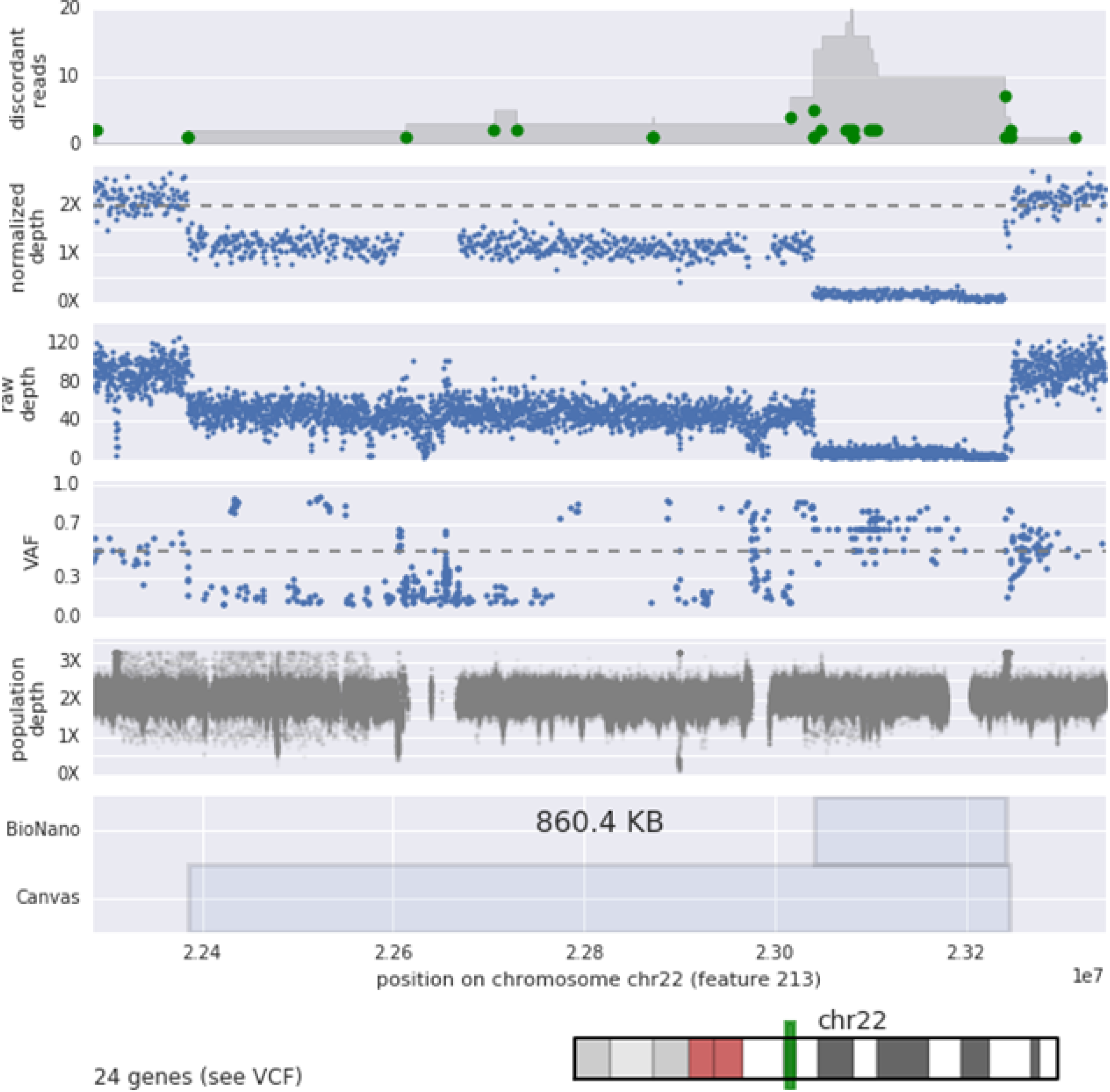
This is a homozygous deletion flanking a mosaic 22q11 deletion, a likely cell line artifact. Our WGS pipeline called this event as a single CNV, whereas the BioNano/PacBio caller only called the homozygous deletion.

**Supplemental Note Figure 2:**
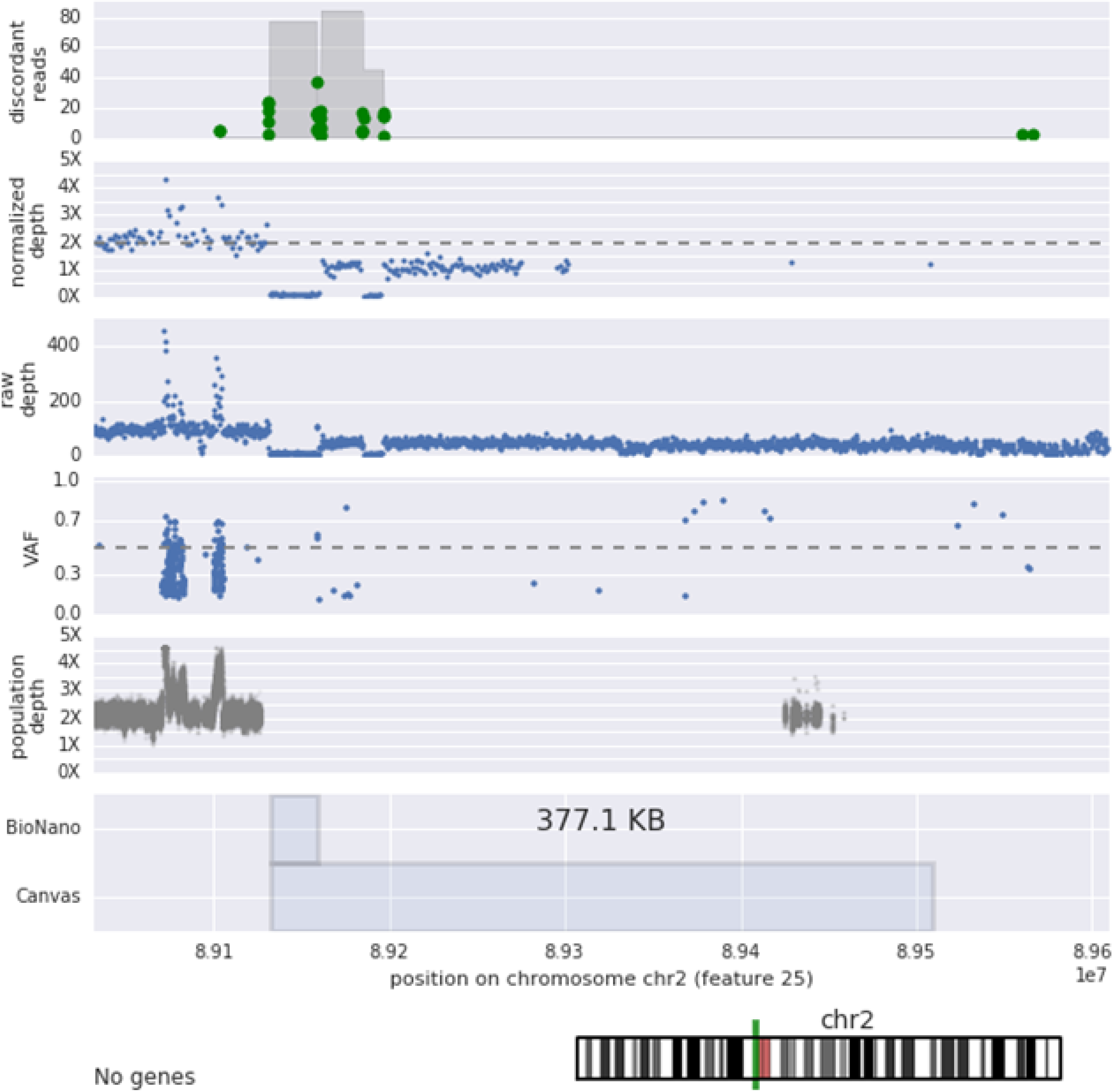
This CNV is a homozygous deletion followed by a mosaic loss leading up to the centromere of chromosome 2. The homozygous deletion is called by BioNano, but the mosaic loss is missed or filtered.

**Supplemental Note Figure 3:**
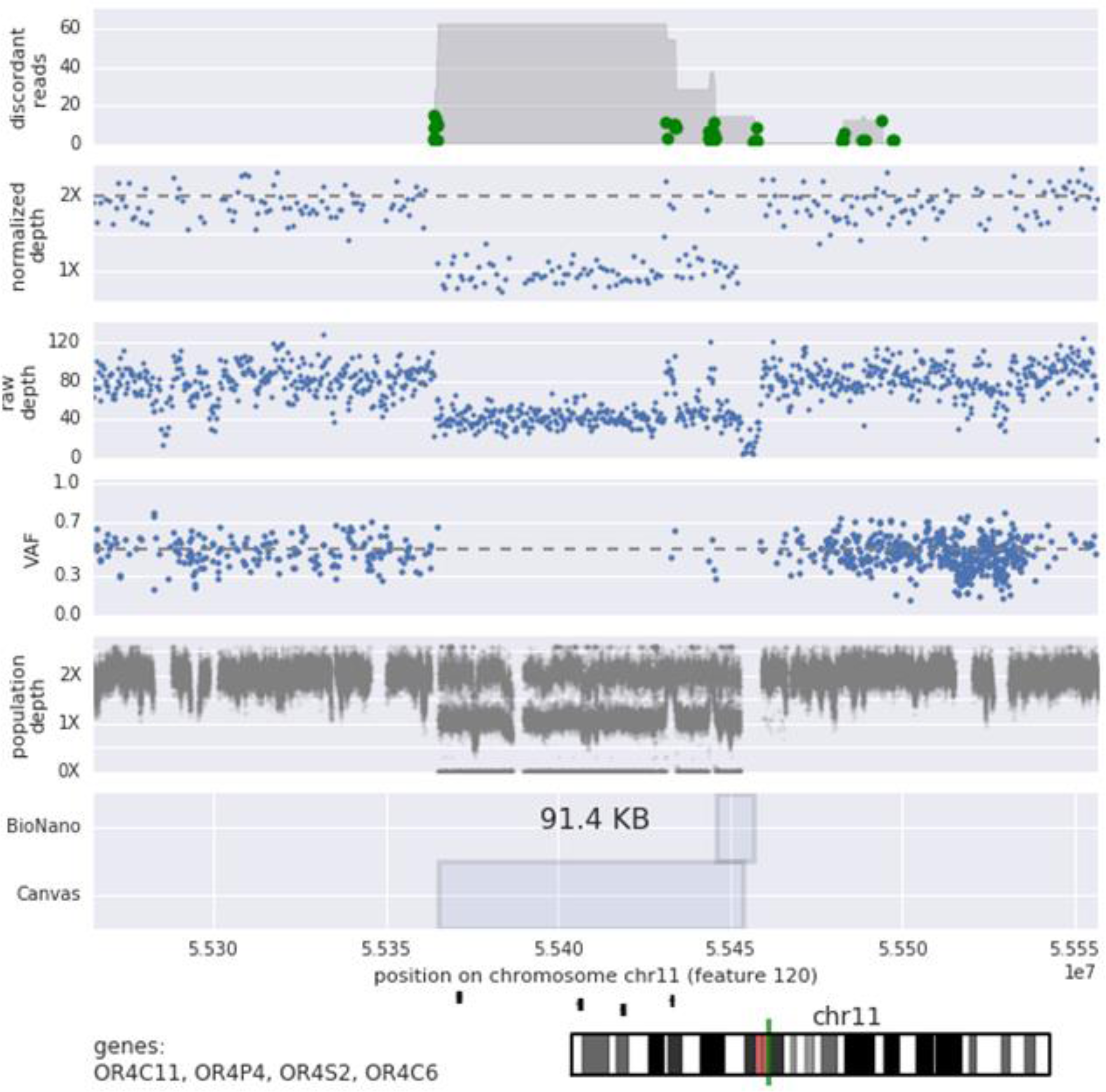
This is a very common deletion supported by both population data, as well as discordant sequencing reads. BioNano only partially called this deletion, but the data strongly support the WGS depth based call. We suspect that this was missed by BioNano due to the presence of more complex structural rearrangement in the region.

## Supplemental Note: Manual Inspection of Coriell CNV Calls

We conducted an investigation of false negative (FN) calls to determine if any systematic issues could be identified. To search for error modes, FN calls were analyzed via manual inspection of microarray depth, sequencing depth, and discordant reads. We found nearly all of the discrepant calls occurred in low complexity regions not covered by microarray, or had ambiguous annotation on the Coriell website and/or copy number calling publication (Tang et al., 2013). Although we cannot definitively conclude that certain calls from Coriell are erroneous, data from NGS and multiple genotyping arrays do not support a majority of these calls. To this effect, while the initial recall was calculated at 86% (31/36) events, this in-depth view of data leads us to speculate that the sensitivity is considerably higher.

### Manual Inspection of Coriell CNV Calls with Disagreement of Boundaries

Prior to validation, a 75% reciprocal overlap threshold was set for calling of concordant calls. In **Table 1** we note 4 CNV calls with reciprocal overlaps in the range of 50-75%. A post-hoc analysis of this data generally support the boundaries of the Canvas CNV. The Coriell provided coordinates for all four CNVs are provided in Table S1.

NA02767: trisomy 21. The Coriell website records the CNV as extending across the centromere, whereas canvas calls the trisomy as the entirety of 21q, resulting in a 70% overlap. We note that we cannot call CNVs into the centromeres due to high sequence complexity.

NA06804: HPRT1 duplication. The Coriell website reports a qualitative description of exon 2 and 3 duplication. The Canvas call is shifted from the ‘truth-set’ coordinates by 3kb, but is well supported by both the read depth, as well as discordant read data.

**Supplemental Note Figure 4:**
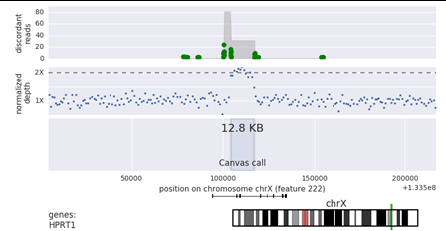
Depth and discordant read locations in Coriell sample NA06804 near the HPRT1 locus.

NA11428: 3q duplication. This large copy number duplication was split into two calls due to the presence of a 5kb common deletion in one of the genomic DNA copies. The segmentation resulted in the GAIN to be split into two large CNV calls comprising 44% and 52% of the truth set duplication. We note that such segmentation is common for large CNVs and protocols are in place for the ICSL clinical workflow to address and merge such calls (Methods).

**Supplemental Note Figure 5:**
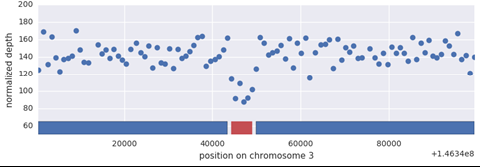
Depth bins visualized for Coriell sample NA11428. The blue and red bars at the bottom of the figure indicate the results of the CNV partitioning.

### Manual Inspection of False Negative Coriell CNV Calls

Independent investigation of false negative (FN) calls was performed to determine if any systematic issues could be identified (Supplemental Note Table 1). To search for error modes, FN calls were analyzed via manual inspection of microarray depth, sequencing depth, and discordant reads.

**Supplemental Note Table 1:** Investigation of false negative Coriell CNV calls.

| Coriell Call | Subject | Chrom | Size | Coriell CN | Array evidence | Manual inspection evidence | Suspected reason for FP |
| --- | --- | --- | --- | --- | --- | --- | --- |
| 6 | NA06804 | chrX | 13 | 0 | No | No | Misinterpretation of primary data |
| 7 | NA09834 | chr19 | 105 | 1 | No | No | Poor mapping or array artifact |
| 30 | NA20304 | chr15 | 418 | 1 | No | No | Poor mapping or array artifact |
| 33 | NA21886 | chr2 | 102 | 3 | No | No | Bad annotation in Coriell |
| 37 | NA21886 | chr4 | 112 | 1 | No | No | Bad annotation in Coriell |

Five CNVs displayed ambiguous annotation on the Coriell web site or associated publication (Tang et al., 2013), and only a single event was replicated in the Canvas call-set, while the remaining events had very little support from any data source. The resulting discrepancies indicate that the Coriell calls could be the results of artifacts in the experimental or bioinformatics analysis of these cell lines, or could be events originating at the cell line level that have diverged between different cell line specimens. For an example of a call with no array support see **Supplemental Note Figure 6**.

One false negative from the Coriell call-set call occurred in the NA6804 sample on the first intron of the HPRT1 gene. Re-inspection of the literature supporting this event showed conflicting reports of this pathogenic rearrangement. Yang et al. (Yang et al., 1984, 1988) report a duplication of exons 2 and 3 of the gene alongside a deletion of exon 1. In contrast Monnat et al. (Monnat et al., 1992) showed that the gain in exons 2/3 results from an insertion of the sequence into the first intron of HPRT1. While Canvas correctly identified the reported duplication, the read depth and paired read data seem to support the latter report of an insertion of this sequence into the first intron (**Supplemental Note Figure 4**). Based off of this evidence as well as Monnet et al. it is likely that the reported deletion is actually an artifact of the experimental methodology of Yang et al. as opposed to a true CNV.

Another potential false-positive call was a 418kb deletion on chromosome 15 in NA20304 (**Supplemental Note Figure 7**). While we are able to observe this deletion in the raw sequencing data, we note that this region contains highly redundant genomic sequence, which caused the Canvas caller to be unable to assign a normalized sequencing depth to this region. We also note very few probes in this region for both the 850k and 2.5M Illumina microarrays reflecting the likely due to inability to construct unique primers in this region. Taken together we hypothesize that this CNV could be an artifact of the Affymetrix array from which it was derived, but have insufficient evidence to definitively rule this call out as a false negative.

**Supplemental Note Figure 6:**
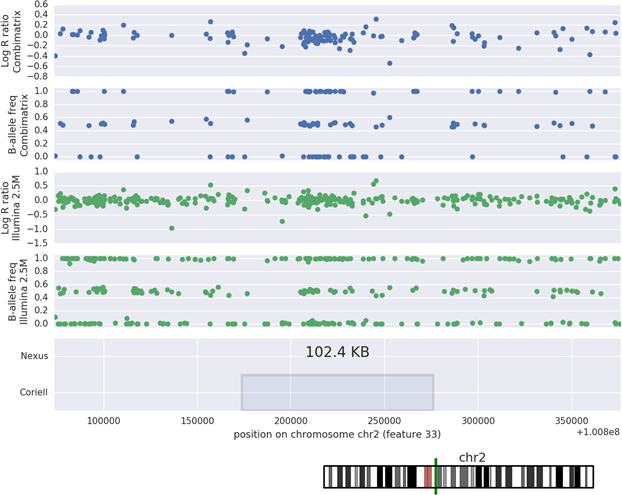
Example of CNV annotated in the Coriell sample NA21886 that has little to no support from two commonly used clinical microarrays.

**Supplemental Note Figure 7:**
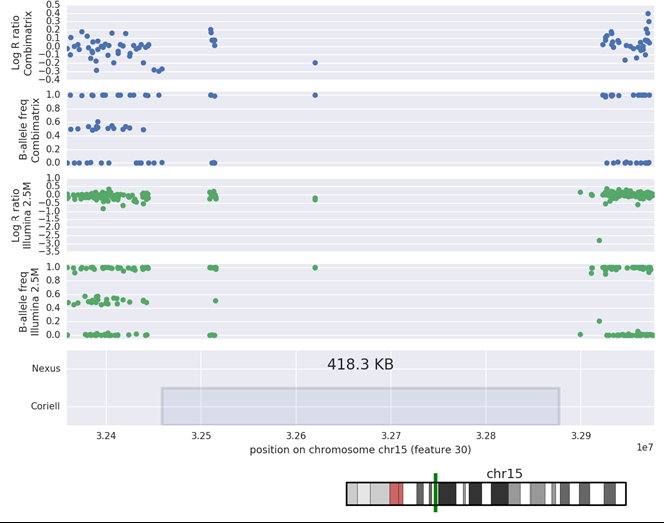
Example of CNV annotated in the Coriell sample NA20304 that has little to no support from two commonly used clinical microarrays.

## Supplemental Note: *de novo* CNV phasing models

For de-novo CNVs we observe the inheritance patterns of small variants to decipher parental haplotype on which a CNV resides.

### Deletion phasing

Here we simply compare inheritance of variants under the assumption of the deletion being on either the maternal or paternal alleles.

**Supplemental Note Table 2:**
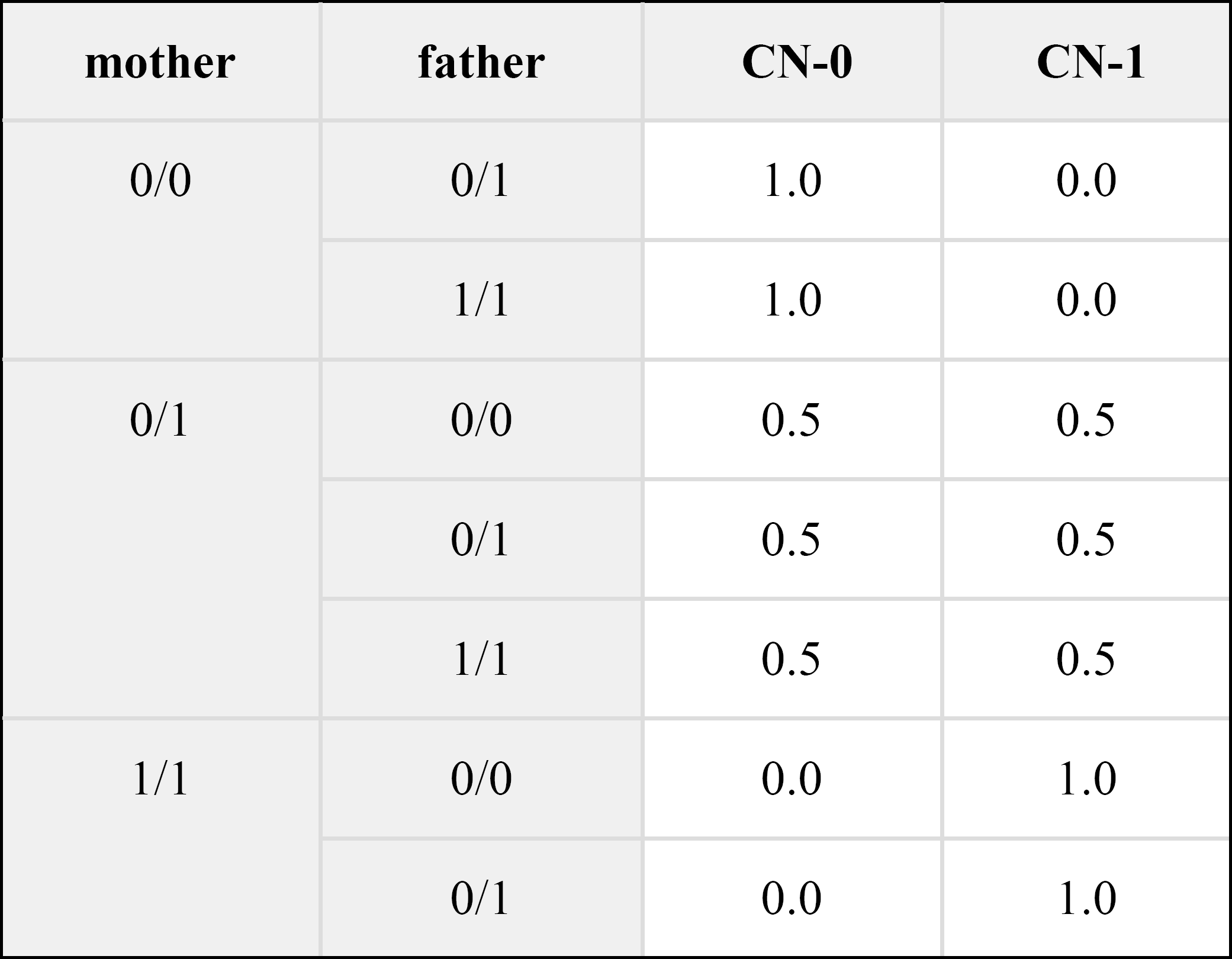
Model assuming deletion on **paternal** allele (all variants inherited from mother):

| **mother** | **father** | **CN-0** | **CN-1** |
| --- | --- | --- | --- |
| 0/0 | 0/1 | 1.0 | 0.0 |
|  | 1/1 | 1.0 | 0.0 |
| 0/1 | 0/0 | 0.5 | 0.5 |
|  | 0/1 | 0.5 | 0.5 |
|  | 1/1 | 0.5 | 0.5 |
| 1/1 | 0/0 | 0.0 | 1.0 |
|  | 0/1 | 0.0 | 1.0 |

#### Example deletion

3MB deletion on the paternal allele. See Supplemental Note Figure 8 for illustration of model transition frequencies.

#### Model log-likelihoods

father -2472.071702

mother -6295.203394

**Prediction:** *de novo* deletion on paternal allele.

**Supplemental Note Figure 8:**
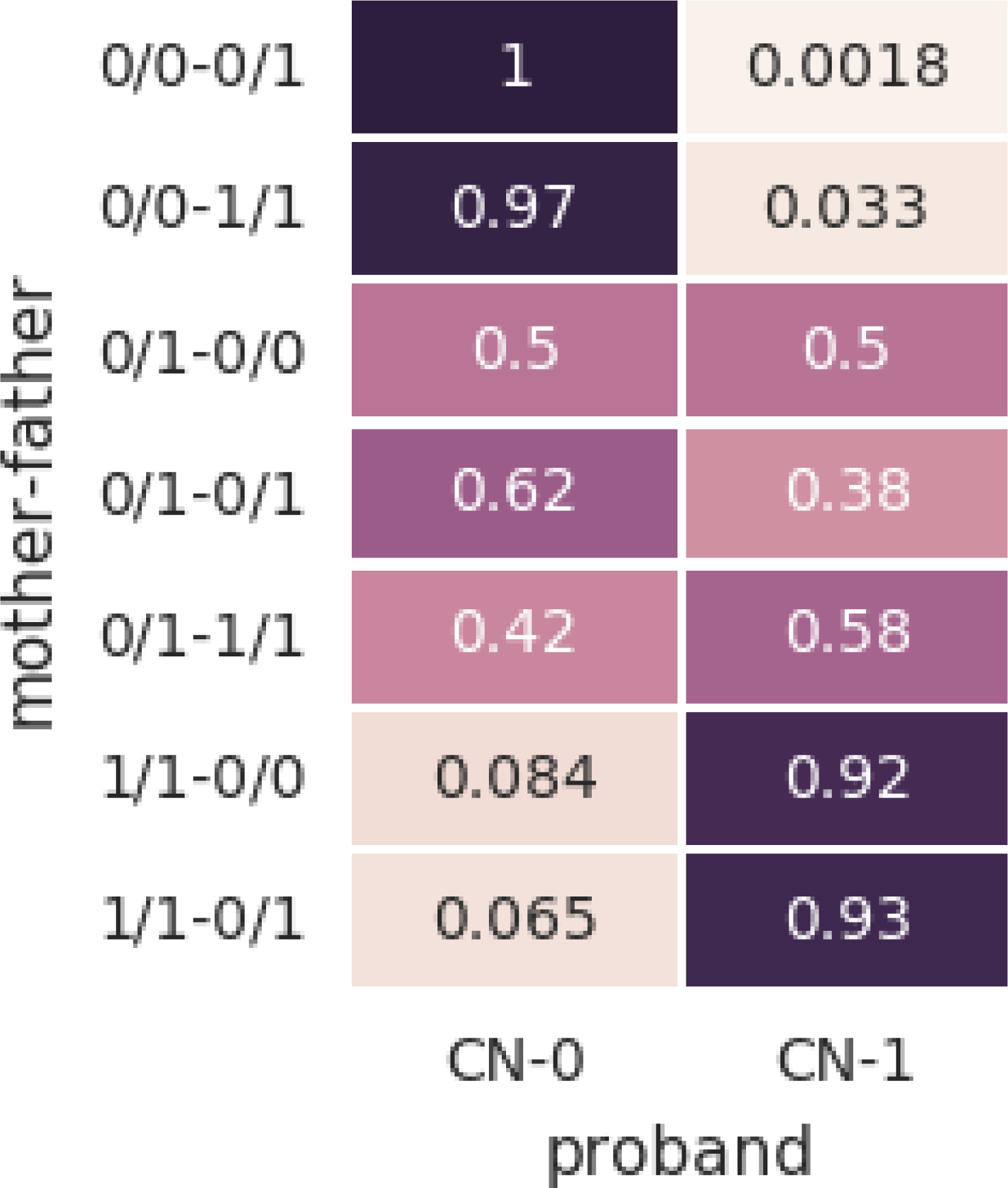
Transition frequencies for example deletion.

### Gain phasing

For gains, there are four possible scenarios. A gain may be of maternal or paternal origin, and be either simple or complex. By simple we refer to a duplication of a single allele, while a complex gain refers to the scenario where a proband can inherit material from both parents’ copies of the DNA segment (an example of this is in an unbalanced translocation).

Additionally, rather than having two copy states as in the case of deletions, gains have four possible variant copy states.

**Supplemental Note Table 3:** Model assuming a **simple duplication** of a **maternal** allele:

| mother | father | CN-0 | CN-1 | CN-2 | CN-3 |
| --- | --- | --- | --- | --- | --- |
| 0/0 | 0/1 | 0.50 | 0.50 | 0.00 | 0.00 |
|  | 1/1 | 0.00 | 1.00 | 0.00 | 0.00 |
| 0/1 | 0/0 | 0.50 | 0.00 | 0.50 | 0.00 |
|  | 0/1 | 0.25 | 0.25 | 0.25 | 0.25 |
|  | 1/1 | 0.00 | 0.50 | 0.00 | 0.50 |
| 1/1 | 0/0 | 0.00 | 0.00 | 1.00 | 0.00 |
|  | 0/1 | 0.00 | 0.00 | 0.50 | 0.50 |

**Supplemental Note Table 4:** Model assuming inheritance of **both maternal alleles** (along with one paternal allele):

| mother | father | CN-0 | CN-1 | CN-2 | CN-3 |
| --- | --- | --- | --- | --- | --- |
| 0/0 | 0/1 | 0.5 | 0.5 | 0.0 | 0.0 |
|  | 1/1 | 0.0 | 1.0 | 0.0 | 0.0 |
| 0/1 | 0/0 | 0.0 | 1.0 | 0.0 | 0.0 |
|  | 0/1 | 0.0 | 0.5 | 0.5 | 0.0 |
|  | 1/1 | 0.0 | 0.0 | 1.0 | 0.0 |
| 1/1 | 0/0 | 0.0 | 0.0 | 1.0 | 0.0 |
|  | 0/1 | 0.0 | 0.0 | 0.5 | 0.5 |

#### Example gain

7MB deletion on the paternal allele.

Allele fraction across genotypes clearly shows the dependence of copy number state on parental genotypes (**Supplemental Note Figure 9**).

#### Model likelihoods

father-complex -7948

father-dup -26755

mother-complex -19933

mother-dup -24622

**Prediction:** The gain is resultant from inheritance of both paternal alleles, along with a single maternal allele.

**Supplemental Note Figure 9:**
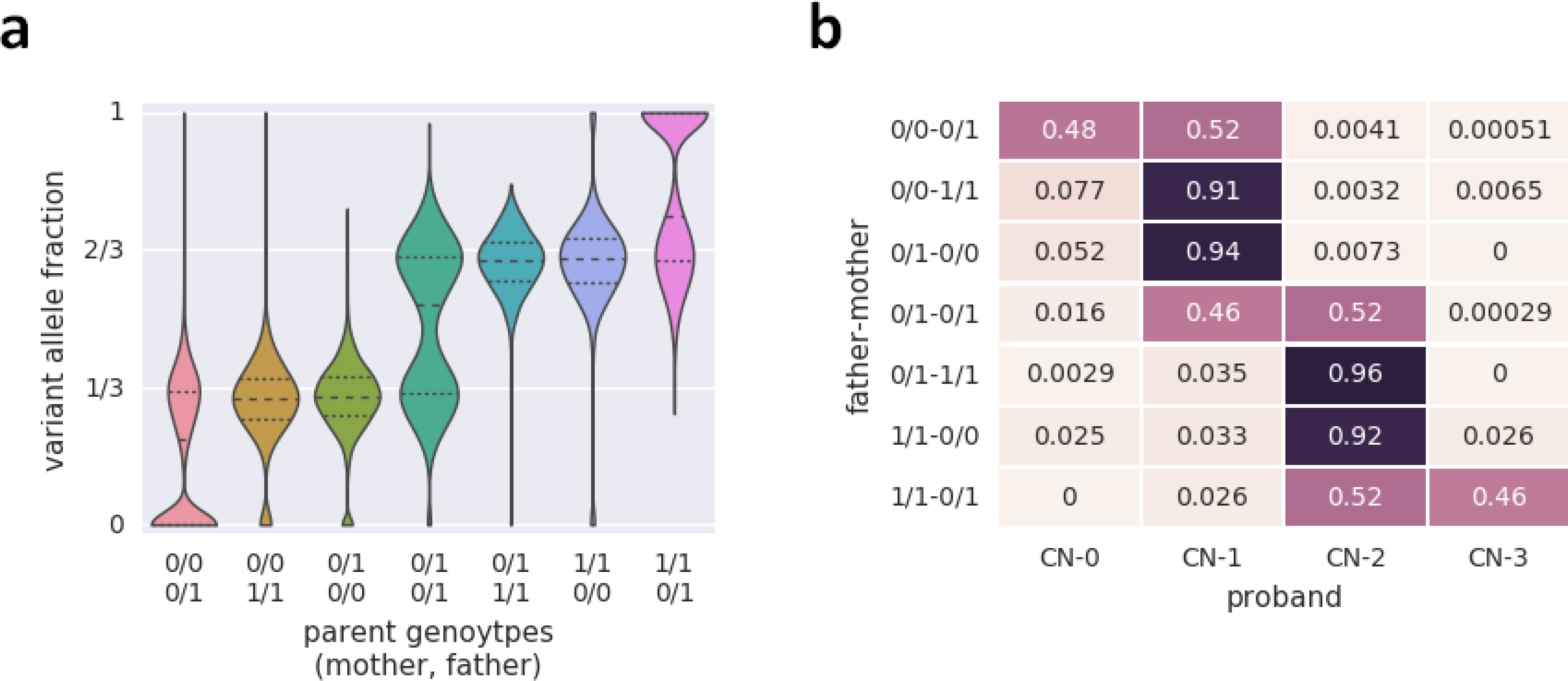
Variant allele fraction across parental genotypes (a) and copy-number state transition frequencies (b) for example duplication.

